# Correction of respiratory artifacts in MRI head motion estimates

**DOI:** 10.1101/337360

**Authors:** Damien A. Fair, Oscar Miranda-Dominguez, Abraham Z. Snyder, Anders Perrone, Eric A. Earl, Andrew N. Van, Jonathan M. Koller, Eric Feczko, Rachel L. Klein, Amy E. Mirro, Jacqueline M. Hampton, Babatunde Adeyemo, Timothy O. Laumann, Caterina Gratton, Deanna J. Greene, Bradley L. Schlaggar, Don Hagler, Richard Watts, Hugh Garavan, Deanna M. Barch, Joel T. Nigg, Steven E. Petersen, Anders Dale, Sarah W. Feldstein-Ewing, Bonnie J. Nagel, Nico U.F. Dosenbach

## Abstract

Head motion represents one of the greatest technical obstacles for brain MRI. Accurate detection of artifacts induced by head motion requires precise estimation of movement. However, this estimation may be corrupted by factitious effects owing to main field fluctuations generated by body motion. In the current report, we examine head motion estimation in multiband resting state functional connectivity MRI (rs-fcMRI) data from the Adolescent Brain and Cognitive Development (ABCD) Study and a comparison ‘single-shot’ dataset from Oregon Health & Science University. We show unequivocally that respirations contaminate movement estimates in functional MRI and that respiration generates apparent head motion not associated with degraded quality of functional MRI. We have developed a novel approach using a band-stop filter that accurately removes these respiratory effects. Subsequently, we demonstrate that utilizing this filter improves post-processing data quality. Lastly, we demonstrate the real-time implementation of motion estimate filtering in our FIRMM (Framewise Integrated Real-Time MRI Monitoring) software package.

## Introduction

Head motion represents one of the greatest technical obstacles in human magnetic resonance imaging (MRI). In both task-driven fMRI (t-fMRI) and resting state functional connectivity MRI (rs-fcMRI), even sub-millimeter head movements (i.e., micro-movements) introduce artifact, thereby corrupting the data in both typical and atypical populations (Fair et al., 2012; Power et al., 2012; Satterthwaite et al., 2012; K. R. Van Dijk et al., 2012; Yan et al., 2013). Consequently, much effort has been devoted to the development of post-acquisition artifact reduction methods (Behzadi et al., 2007; Burgess et al., 2016; Ciric et al., 2017; Di Martino et al., 2014; Griffanti et al., 2014; Jo et al., 2013; Kundu et al., 2013; Muschelli et al., 2014; Patel et al., 2014; Power et al., 2013, 2012; Power, 2016; Pruim et al., 2015; Salimi-Khorshidi et al., 2014; Satterthwaite et al., 2012, 2013; Joshua S. Siegel et al., 2016; K. R. a Van Dijk et al., 2012). However, while these efforts have been helpful in reducing motion artifact, many come at a cost – namely, the risk of losing entire subjects after discovering insufficient amounts of quality data exist for a given participant.

To address these difficulties, we developed <u>F</u>ramewise <u>I</u>ntegrated <u>R</u>eal-time <u>M</u>RI <u>M</u>onitoring (FIRMM) (Dosenbach et al., 2017). FIRMM provides near instantaneous analysis of head motion (framewise displacement = FD) during scanning. This feature enables the duration of resting state fMRI scanning to be dynamically determined, that is, for as long as is necessary to acquire a prescribed quantity of data meeting a fixed quality assurance criterion. Thus, FIRMM helps to ensure that an adequate quantity of resting state data is acquired in most participants.

It also reduces the need for ‘overscanning’ all participants in order to ensure adequate acquisitions across subjects. Moreover, subjects who cannot suppress excessive motion can be identified promptly and efficiently excluded from the study. Additionally, FIRMM alerts the scanner operator to changes in participant behavior (e.g., sleep, or increased movement because of discomfort), and also can be utilized to give motion feedback to participants themselves. These advances have been thoroughly detailed in prior reports (Dosenbach et al., 2017; Greene et al., 2018).

Here, we address new issues that have arisen when calculating head motion estimates (FD) fMRI data. Multiband imaging (simultaneous multi-slice sequences [SMS]) is a recently developed technique that substantially speeds up fMRI by exciting multiple slices simultaneously (Feinberg and Yacoub, 2012; Moeller et al., 2010; Uğurbil et al., 2013; Xu et al., 2013). Thus, the time needed to acquire one whole brain volume can be reduced to well below 1 sec. While multiband sequences offer several clear advantages and are now becoming widely used, they come at the cost of reducing the temporal signal to noise ratio (tSNR; (Chen et al., 2015)) and, potentially, new types of ‘slice-leakage’ artifacts (Barth et al., 2016; Todd et al., 2016).

Another unanticipated consequence of the improved temporal resolution of SMS sequences is corruption of head motion estimation (FD) arising from the interaction between echo-planar imaging (EPI) and small perturbations of the main (B0) field generated by changes in body position. For example, given a bandwidth of 2646 Hz/pixel in the phase-encoding direction and 2.4 mm pixels (as in the ABCD study), a B0 perturbation of 15 parts per million (15*123 = 1845 Hz in k-space) will lead to a 1.67 mm shift of the reconstructed image in that direction. Such shifts are easily corrected during retrospective rigid body motion correction; however, and critically, there may be no associated true head motion and, therefore, no artefactual changes in blood oxygen level dependent (BOLD) signals on the basis of spin history effects (Friston et al., 1996a). It follows that factitious head motion should not be taken as an indicator of degraded image quality.

The most obvious candidate for a process that might modulate B0 to generate factitious estimates in our motion estimates is chest movement from breathing. It has long been recognized that respirations change the magnetic field (B0)(Van de Moortele et al., 2002), and SMS sequences might be exacerbating these changes. Investigators using SMS sequences tend to utilize higher sampling frequencies. As noted above, these higher sampling frequencies (shorter time-to-repeat [TR]) might also increase the prominence of respiratory B0 modulation in the motion traces. Several of our most prominent multi-site functional MRI studies employ SMS approaches (The Human Connectome Project, The Adolescent Brain and Cognitive Development Study, etc.). Several researchers have remarked that higher sampling frequencies seem to unmask the effects of respirations on head motion estimates (Joshua S Siegel et al., 2016). Importantly, such effects might also have an unintended consequence of affecting the quality of structural scanning of motionless participants when using volumetric navigators (vNAVs) (Tisdall et al., 2016; Zaitsev et al., 2017). However, the effects of respiration on head motion estimates, and subsequently, methodologies to correct factitious motion estimates have not been thoroughly studied.

In the current report, we examine the effects of respirations on head motion estimates using data from the Adolescent Brain and Cognitive Development (ABCD) Study, and a separate, inhouse ‘single-shot’ dataset. We show unequivocally that respirations inflate movement estimates. We then present a novel approach that utilizes a band-stop (or notch) filter to remove respiration-related effects from the motion estimates. Subsequently, we emphasize that utilizing such a filter improves data quality and outcomes. Lastly, we demonstrate an implementation of our filter in real-time to improve the accuracy of real-time estimates of brain motion using FIRMM.

## Materials and Methods

### Participants

#### ABCD – Multiband Dataset

For the current report, we analyzed data acquired in the subset of ABCD participants (see (Bjork et al., 2017; Casey et al., 2018; Lisdahl et al., 2018; Volkow et al., 2017) scanned at Oregon Health & Science University (OHSU). The ABCD study targets child participants that are ethnically and demographically representative of United States (US) population. Participants and their families are recruited through school- and community-based mailings in 21 centers across the US, following locally and centrally approved Institutional Review Board approved procedures. Written informed consent is obtained from parents, and written assent obtained from children. All ABCD participants are 9-11 years of age at study entry. Exclusion criteria include current diagnosis of a psychotic disorder (e.g., schizophrenia), a moderate to severe autism spectrum disorder, intellectual disability, or alcohol/substance use disorder, lack of fluency in English (for the child only), uncorrectable sensory deficits, major neurological disorders (e.g., cerebral palsy, brain tumor, multiple sclerosis, traumatic brain injury with loss of consciousness > 30 minutes), gestational age < 28 week or birth-weight < 1.2kg.), neonatal complications resulting in > 1 month hospitalization following birth, and MRI contraindications (e.g., braces). A total of 62 children are examined here (Female 32; Mean Age = 11.65). All presently analyzed ABCD participants chosen here had both sufficient EPI data (4 × 5 minute runs) and quality physiologic data obtained from Siemens built in physiologic monitor and respiratory belt.

Prior to MRI scanning, respiratory monitoring belts were placed comfortably around the child’s ribs (with sensor horizontally aligned just below the ribcage). A pulse oxygen monitor was placed on the non-dominant index finger. Participants were scanned on a Siemens 3.0 T Magnetom Prisma system (Siemens Medical Solutions, Erlangen, Germany) with a 32-channel head coil, located at OHSU’s Advanced Imaging Research Center. A high-resolution T1-weighted MPRAGE sequence was acquired (resolution = 1 × 1 × 1 mm). BOLD-weighted functional images were collected (along the anterior–posterior commissure) using T2*-weighted echo planar imaging (TR = 0.80 ms, TE = 30 ms, flip angle = 52, FOV = 216 mm2, 60 slices covering the entire brain, slice thickness = 2.4 mm, resolution = 2.4 × 2.4 × 2.4 mm, MB acceleration = 6). Further details can be seen in Casey et al. (Casey et al., 2018). Four runs of 5 min of resting state BOLD data were acquired, during which participants were instructed to stay still and fixate on a white crosshair in the center of a black screen projected from the head of the scanner and viewed with a mirror mounted on the 32-channel head coil. One additional research volunteer (male, aged 25) underwent the same protocol with the TR variably at 0.8s, 1.5s, and 2.0s. This same participant also underwent a non-multiband sequence similar to the OHSU data set noted below (TR = 2500 ms, TE = 30 ms, flip angle = 90, FOV = 240 mm^2^, 36 slices covering the entire brain, slice thickness = 3.8 mm, resolution = 3.75 × 3.75 × 3.8 mm).

#### OHSU – Single Band Dataset

Single band data were acquired in 321 total scanning sessions (177 female, Mean Age = 11.16) right-handed children over 416 scanning sessions. These data were obtained at the Advanced Imaging Research Center at OHSU during the course of ongoing longitudinal studies (Costa Dias et al., 2015; Dosenbach et al., 2017; Feczko et al., 2017; Gates et al., 2014; Grayson et al., 2014; Sarah L Karalunas et al., 2014; Mills et al., 2017; Kathryn L Mills et al., 2012; Miranda-Dominguez et al., 2017; Ray et al., 2014). Children were recruited by mass mailings to the local community. Only neurotypical controls contributed to the present results. Their diagnostic status—that is, confirmation of typically developing control children without major psychiatric, behavioral, or developmental disorder, was carefully evaluated with a multi-informant, multimethod research clinical process. This process included nationally normed standardized rating scales, semi-structured clinical interviews, and expert review (for more details see (Costa Dias et al., 2015; Fair et al., 2010; Sarah L. Karalunas et al., 2014; Kathryn L. Mills et al., 2012; Nigg et al., 2018)). For this report, exclusion criteria were ADHD, tic disorder, psychotic disorder, bipolar disorder, autism spectrum disorder, conduct disorder, major depressive disorder, intellectual disability, neurological illness, chronic medical problems, sensorimotor disability, significant head trauma (with loss of consciousness), or any concurrent psychotropic medication. Children were also excluded if they had contraindications to MRI. The Human Investigation Review Board at OHSU approved the research. Written informed consent was obtained from respective parents and verbal or written assent was obtained from child participants.

Participants were scanned on a Siemens Tim Trio 3.0 T Magnetom Tim Trio system (Siemens Medical Solutions, Erlangen, Germany) with a 12-channel head coil. Visual stimuli were viewed via a mirror mounted on the head coil. One high-resolution T1-weighted MPRAGE sequence was acquired (resolution = 1 × 1 × 1 mm). BOLD-weighted functional images were collected using T2*-weighted echo planar imaging (TR = 2500 ms, TE = 30 ms, flip angle = 90, FOV = 240 mm^2^, 36 slices covering the entire brain, slice thickness = 3.8 mm, resolution = 3.75 × 3.75 × 3.8 mm). Three resting state runs of duration 5 min were acquired during which participants were instructed to stay still and fixate on a white crosshair in the center of a black screen.

#### Post-acquisition Processing

All data were processed following slightly modified pipelines developed by the Human Connectome Project (Glasser et al., 2013; Mills et al., 2017; Miranda-Dominguez et al., 2017). These pipelines require the use of FSL (Jenkinson et al., 2012; Stephen M Smith et al., 2004; Woolrich et al., 2009) and FreeSurfer (Dale et al., 1999; Fischl, 2012; Fischl and Dale, 2000). Since T2-weighted images were not acquired in all individuals this aspect of the pipeline was omitted. Gain field distortion corrected T1-weighted volumes were first aligned to the MNI AC-PC axis and then non-linearly normalized to the MNI atlas. Later, the T1-weighted (T1w) volume was re-registered to the MNI template (Fonov et al., 2011) using boundary based registration (Greve and Fischl, 2009) and segmented using the recon-all procedure in FreeSurfer. The BOLD data were corrected for magnetization inhomogeneity-related distortions using the TOPUP module in FSL (Stephen M. Smith et al., 2004). An average volume was calculated following preliminary rigid body registration of all frames to the first frame. This average volume was registered to the T 1w. Following composition of transforms, all frames were resampled in register with T1-weighted volume in a single step.

##### Surface registration

The cortical ribbon was segmented out of the T1-weighted volume and represented as a tessellation of vertices. The co-registered BOLD data then were projected on these vertices. Vertices with a high coefficient of temporal variation, usually attributable to susceptibility voids or proximity to blood vessels, were excluded. Next, the vertex-wise time series were down-sampled to a standard surface space (grayordinates) and geodesically smoothed using a 2mm full-width-half-max Gaussian filter.

##### Nuisance regression

Minimally processed timecourses generated by the HCP pipeline (Glasser et al., 2013) were further preprocessed to reduce spurious variance, first by regression of nuisance waveforms. Nuisance regressors included the signal averaged over gray matter as well as regions in white matter and the ventricles (Burgess et al., 2016). 24-parameter Volterra expansion regressors were derived from retrospective head motion correction (Friston et al., 1996b; Power et al., 2014, 2012). Nuisance regressor beta weights were calculated solely on the basis of frames with low movement but regression was applied to all frames in order to preserve the temporal structure of the data prior to filtering in the time domain. Timecourses then were filtered using a first order Butterworth band pass filter retaining frequencies between 0.009 and 0.080 Hz.

#### Rigid body motion correction

Each data frame (volume) is aligned to the first frame of the run through a series of rigid body transform matrices, *T_i_*, where *i* indexes frame and the reference frame is indexed by 0. We use the same approach for both real-time and post-processing estimates. Each transform is calculated by minimizing the registration error,

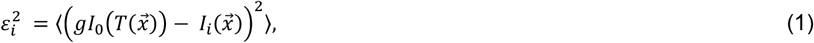

where 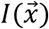 is the image intensity at locus 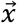 and *g* is a scalar factor that compensates for fluctuations in mean signal intensity, spatially averaged over the whole brain (angle brackets). Each transform is represented by a combination of rotations and displacements. Thus,

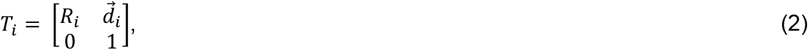

where *R_i_* is a 3 × 3 rotation matrix and 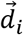 is 3 × 1 column vector of displacements (in mm). *R_i_* is the product of three elementary rotations about the cardinal axes. Thus, *R_i_* = *R_i∝_R_iβ_R_iγ_*, where

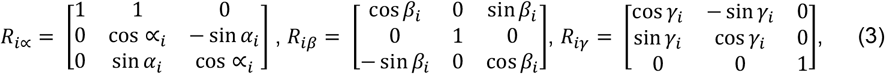

and all angles are expressed in radians.

#### Computation of framewise displacement (FD)

Retrospective head motion correction generates a 6-parameter representation of the head trajectory, *T_i_*, where *i* indexes frame. Instantaneous frame displacement (*FD_i_*) is evaluated as the sum of frame-to-frame change over the 6 rigid body parameters. Thus,

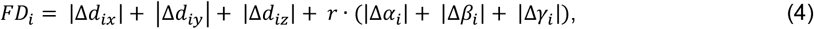

where 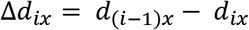, and similarly for the remaining parameters. We assign *r* = 50 mm, which approximates the mean distance from the cerebral cortex to the center of the head for a healthy child or adult. Since *FD_i_* is evaluated by backwards differences, it is defined for all frames except the first.

#### Estimating Respiration characteristics

The nominal rate of respiration changes with age, varying from 30-60 breaths per minute (bpm) at birth to 12-20 bpm at the age of 18 years old. The corresponding frequency in Hz is obtained by dividing bpm by 60. Thus, for example, 20 bpm corresponds to 0.3 Hz. Accordingly, the power spectral density of any signal reflecting respiration should exhibit a peak in the range 0.2 – 0.3 Hz. The width of this peak (line broadening) reflects respiratory rate variability.

It is crucially important to take into account how respiration manifests in BOLD fMRI according to Nyquist theory. The temporal sampling density of BOLD fMRI is the reciprocal of the volume TR. In the ABCD study, the temporal sampling density is 1/(0.8 sec) = 1.25 Hz. The Nyquist folding frequency then is 1.25/2 = 0.625 Hz. This means that signals faster than 0.625 Hz alias to correspondingly lower frequencies. Thus, for example, 0.7 Hz aliases to 0.625 – 0.075 = 0.55 Hz. Importantly, in ABCD data, respiration (0.2 – 0.3 Hz) normally falls well below the Nyquist folding frequency, hence, is not aliased.

In contrast, the BOLD fMRI temporal sampling density in the OHSU dataset (TR = 2.5s) is 1/(2.5s) = 0.4 Hz; the Nyquist folding frequency then is 0.2 Hz. This circumstance suggests that respiration in the OHSU dataset alias into lower bands (these ideas are further tested below). In general, the aliased frequency, *f_a_*, can be calculated as

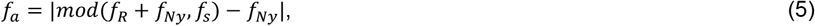

where *f_R_* is the true respiration rate, *f_s_* = 1/*TR* is the fMRI sampling rate and *f_Ny_* = *f_s_*/2 is the Nyquist folding frequency. N.B.: *mod(a,b)* is the modulo function, i.e., the remainder after dividing *a/b*.

#### Estimating motion frequency content

The frequency content of the motion estimates was calculated using power spectral density estimation. To minimize leakage of frequency associated with segmenting the data, we used the standard multitaper power spectral density estimate Matlab function (pmtm) and windowed each segment multiple times using different tapers.

#### Correction of respiration induced artifacts were conducted using a bandstop (notch) filter

Respiration-related signals were removed from the motion estimates using a band-stop (notch) filter. Notch filters remove selected frequency components while leaving the other components unaffected.

A notch filter has 2 design parameters: the central cutoff frequency and the bandwidth (range of frequencies to be eliminated). To determine the central cutoff frequency and the bandwidth for the ABCD participants, we examined the distribution of respiration rates over participants and used the median as cutoff frequency and the quartiles 2 and 3 as bandwidth. Having obtained those values (0.31Hz and 0.43Hz), we used the Second-order IIR notch filter Matlab function, (iirnotch), to design the filter.

#### Post-processing application of bandstop (notch) filter

The designed filter is a difference equation of order 2. Thus, two previous samples are recursively weighted to generate the filtered signal. This procedure starts with the third sample, weights the two previous samples, and continues until the end of the run. Importantly, this filter design introduces phase delay with respect to the original signal. This is a problem because motion estimation then is not in sync with true motion. Phase delay can be eliminated by applying the filter twice, first forward and then backwards.

#### Real-time application of bandstop (notch) filter

Complete elimination of phase delay is not possible in real-time. However, it is possible to minimize phase delay by running the filter in pseudo-real time. Thus, having acquired 5 samples, the filter is applied twice and the best estimate of the filtered signal, free of phase delay, is the result corresponding to the third sample. The process is repeated until the last sample is reached. The filter progressively converges towards the off-line result as more and more data are accumulated. At the end of a given run, the final result is identical to that obtained during post processing.

#### Qualitative assessment was conducted by examining augmented ‘Gray Plots’

Qualitative assessments were conducted by examining the effects of motion on BOLD data using a format introduced by Power *et al.* (Power, 2016). Data from each masked EPI frame is displayed as a vector (representing each voxel; for surface data grayordinates). Each vector, representing each frame in a ‘BOLD run,’ is stacked horizontally in the time domain. This procedure allows all the data from a given run (or full study via concatenated runs) for a given subject to be viewed at once. These rectangular gray plots of BOLD data have been described in detail elsewhere (Power, 2016). Here we make slight modifications. See Figure 3 legend for additional details on how these plots have been augmented for the current report.

#### Quantitative assessment of the impact of factitious head motion

The present objective is to assess the impact of factitious head motion on the quality of BOLD fMRI data and to evaluate alternative filtering strategies for mitigating this impact. Our approach to this problem is based on a procedure introduced by Power et al. (Power et al., 2014) for assessing the link between measured FD and fMRI data quality, originally for the purpose of selecting an optimal frame-censoring threshold. This procedure tests the null hypothesis that measured FD is unrelated to data quality. The involved steps are illustrated in Figure 1. All frames of a given fMRI run are ordered according to increasing FD values. Functional connectivity matrices are computed over sliding windows of 150 volumes (i.e., 1-150, 2-151, …). The mean FC matrix over the first 30 windows (lowest FD) defines a baseline FC matrix. FC matrices are computed over successive windows and their similarity to the baseline is evaluated by correlation of vectorized FC matrices. A relation between FD and fMRI data quality is expected to manifest as a systematic reduction in the correlation between successive windows and the baseline at progressively higher FD values. In the absence of a relation between FD and fMRI data quality, there should be no systematic effect of FD ordering. Thus, the significance of an observed relation can be assessed by comparing the FD-ordered curves to a null distribution generated from surrogate data in which the volumes have been randomly ordered (in our case, 250 permutations). Figure 1 shows a cartoon null distribution of 10 runs for a single subject in black and FD-ordered results in blue. Comparing the true values to the null distribution determines at what FD value the FD-ordered matrices are different from the surrogate data at a given significance (specifically, p < 0.05). In Figure 1, the FD value demarcating the boundary between non-significant (green dot) vs. significant (red dot) FD dependence occurs at about 1.7 mm.

**Figure 1.**
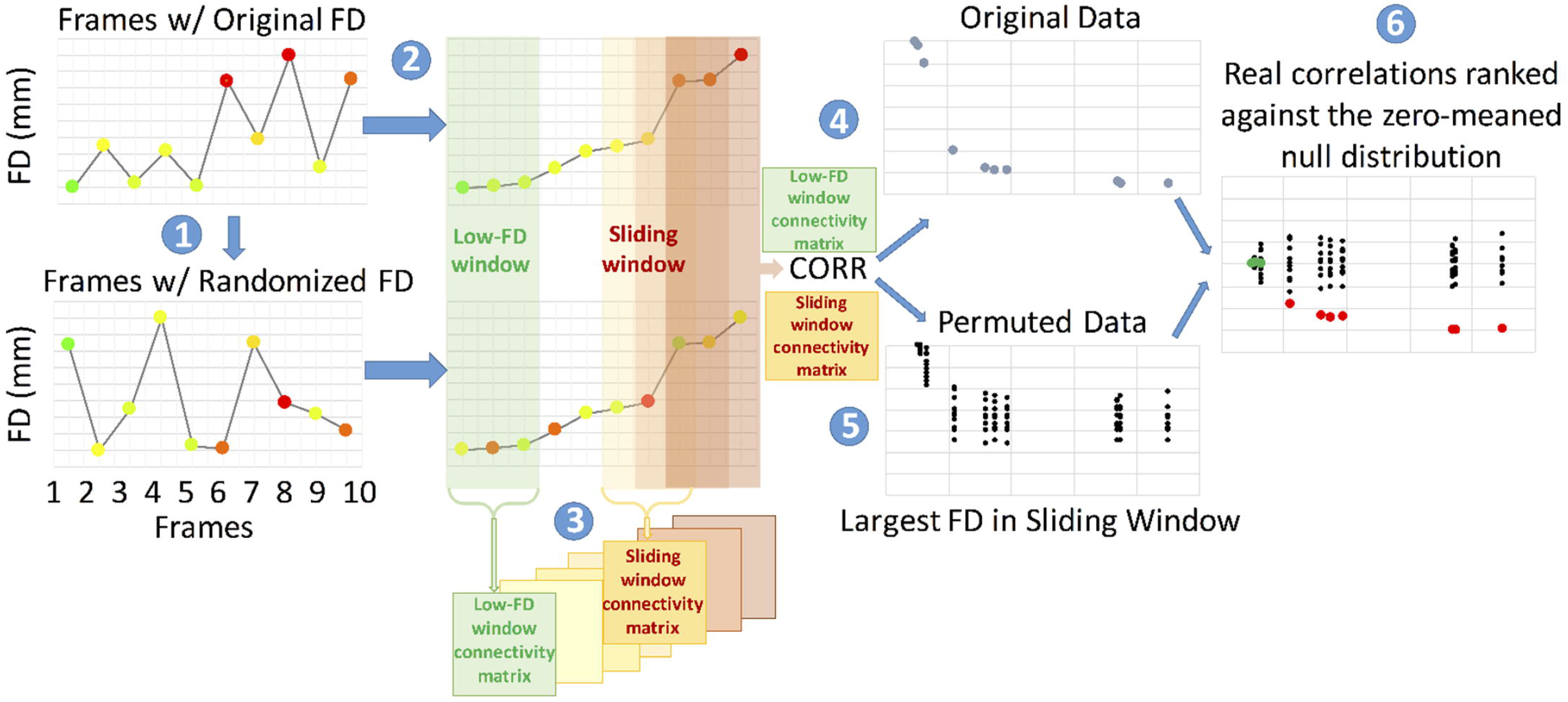
Quantitative assessment of filter success. This approach to quality assessment of the notch filter utilizes all of the data and does not require frame-censoring (Power et al., 2014). The steps for the procedure are outlined in the text and, again, here. Step: (1) All frames of a given fMRI run are ordered according to increasing FD values. (2) Functional connectivity matrices are computed over sliding windows of 150 volumes (i.e., 1-150, 2-151, …). The mean FC matrix over the first 30 windows (lowest FD) defines a baseline FC matrix. (3) FC matrices are computed over successive windows and their similarity to the baseline is evaluated by correlation of vectorized FC matrices. A relation between FD and fMRI data quality is expected to manifest as a systematic reduction in the correlation between successive windows and the baseline at progressively higher FD values. In the absence of a relation between FD and fMRI data quality, there should be no systematic effect of FD ordering. (5) The significance of an observed relation can be assessed by comparing the FD-ordered curves to a null distribution generated from surrogate data in which the volumes have been randomly ordered (in our case, 250 permutations). Values in green are non-significantly different than random. Values in red are significantly different from random. For simplicity of the visualization, the true and null distributions are zero centered.

Figure 2 illustrates how the above-described procedure is used to evaluate alternative strategies for filtering the head motion time series constituting *FD_i_* as defined in Eq. (4). We plot the FD-ordered findings for a single subject against the null (Figure 2A), and then across all subjects (Figure 2B). We then plot the rank of the FD-ordered outcomes across subjects (Figure 2C), and bin these ranks by FD value represented as a heat map (Figure 2D). This procedure provides a vertical distribution of ranks in a FD bin. The cumulative distribution function (CDF) across all FD bins provides an estimate of the average rate at which the windows deviate from the null distribution (Figure 2C,D). If FD is tightly linked to BOLD fMRI data quality, then the shift from randomly distributed ranks occurs at low values of FD. Displacement of this shift towards higher FD values indicates a weaker link between FD and fMRI data quality.

**Figure 2.**
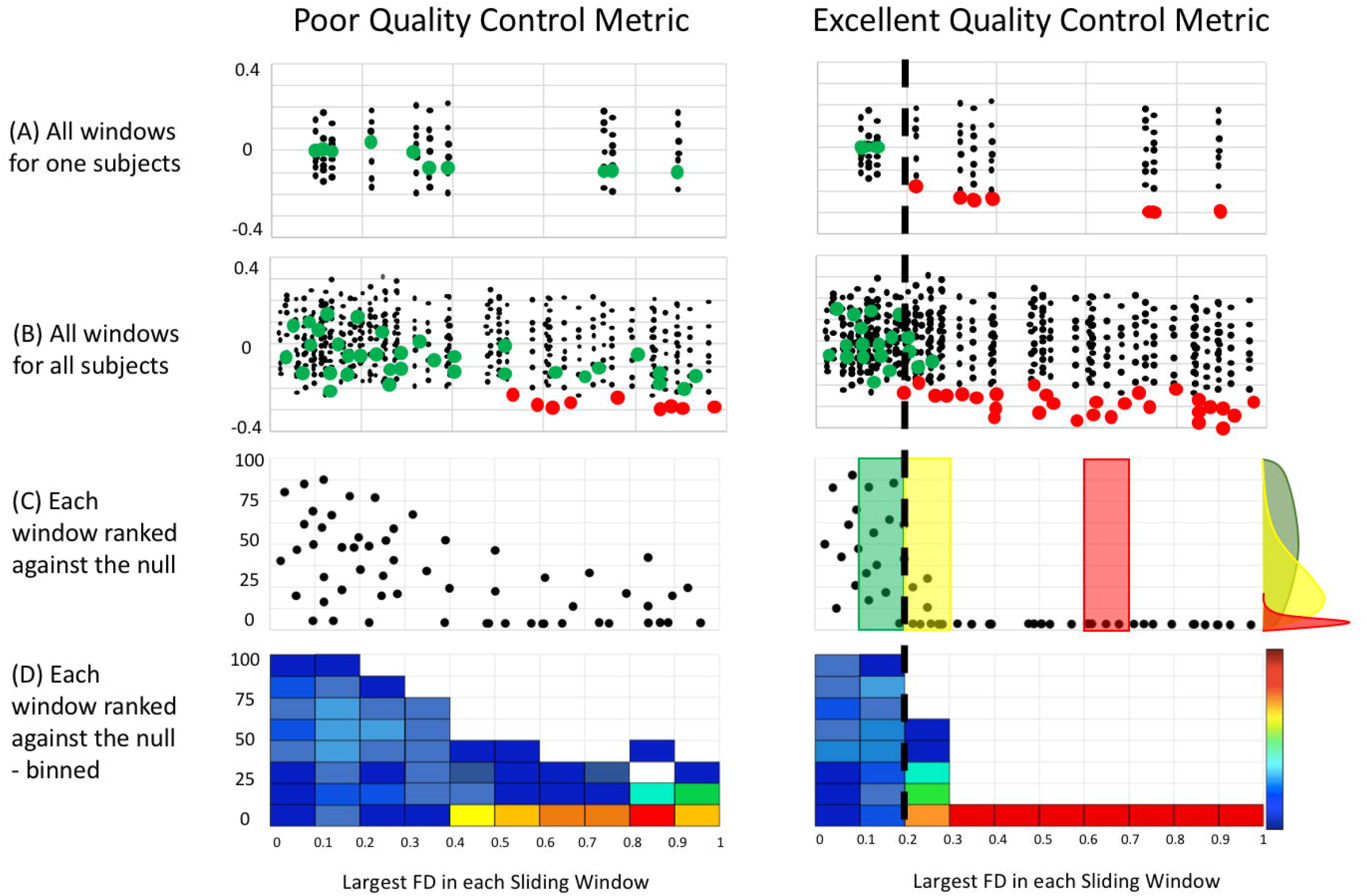
Statistical comparisons between filter conditions. After generating the QC-ordered outcomes of all subjects, and for all conditions (i.e., with and without filter) we can now determine what procedures are the most deviant from random. Here we provide a cartoon demonstrating the procedure. (A) Windows from one participant can be compared for both poor and excellent quality control metrics (e.g. non-filtered and filtered FD measurements) (B) All windows for all participants colored by significance for each filter condition. (C) Percentage rank (relative to null) of the QC-ordered outcomes across subjects, which are binned and represented as a heat map (D). The cumulative distribution function (CDF) across all FD bins provides an estimate of the average rate at which the windows deviate from the null model. The dotted line represents the approximate level at which FD ordered outcomes deviate from random across individuals (i.e., where the distribution of ranks becomes skewed – illustrated with the colored distributions in (C).

## Results

### Fundamental differences in multiband vs. single band motion traces

Figure 3 illustrates the contrast between multiband scanning (TR = 0.8s) vs. single-shot EPI (TR = 2.5s) with regard to the manifestations of respiratory motion in FD traces. The same individual was scanned using both techniques. To clarify the issue at hand, respiration was paced at 0.33Hz by a visual cue in both scans (see SI Movie 1). The gray plots and DVARS traces show that true head motion in this subject was minimal. Nevertheless, in the multiband data, the FD trace frequently rises above 0.2mm and exhibits periodicity corresponding to the imposed respiratory pacing. In comparison, in the single band data, the FD trace generally remains below 0.06mm. These qualitative observations demonstrate a fundamental difference in motion estimates provided by multiband data, as compared to single-shot data.

**Figure 3.**
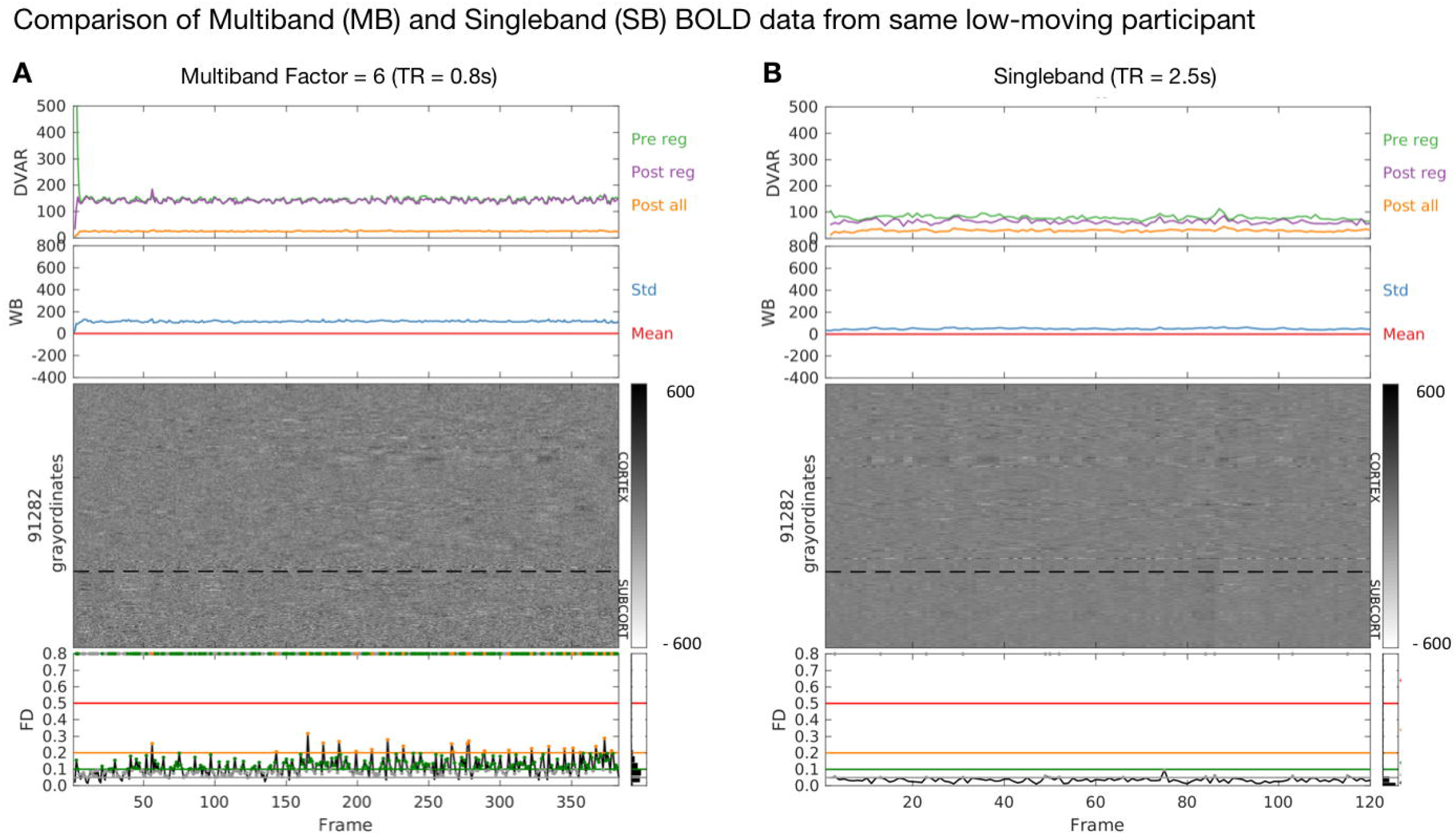
Augmented Gray Plots for a low moving subject (processed data). A) Individual Gray Plot using multiband imaging. B) Gray Plot of same subject with single-shot TR of 2.5 seconds. The gray plots in the middle of the figures represent data from each masked EPI dataset. Each vertically stacked vector represents one frame in a ‘BOLD run.’ Each horizontal vector represents each voxel or surface data grayordinate in the time domain. This procedure allows one to view, in one shot, all the data from a given run (or full study via concatenated runs) for a given subject. On top of the Gray Plots are DVARS measurements (D for derivative of timeseries, VAR for RMS variance across voxels), prior to functional connectivity analysis (green), post the regression phase of the preprocessing (purple), and after the all preprocessing, including filtering (orange). DVARS provides a good estimate of motion that does not rely on retrospective head motion correction (For references, see (Fair et al., 2012; Power et al., 2012; Smyser et al., 2010)). Also included are mean (red) and standard deviation (blue) plots of the whole brain signal (Note: the whole brain signal is flat because these plots represent data after processing, which includes whole brain signal regression). On the bottom of the figures, we plot frame displacement (FD) across the run. Each frame or data point in the FD line plots has a colored circular mark, which represents the FD threshold in which a given frame would have been excluded from future analysis. The corresponding threshold line is displayed by the corresponding color horizontally in the plot. For example, all gray points represent frames that would be excluded at an FD threshold of 0.6 (unless they would also be excluded at a higher threshold), green points represent an FD threshold of 0.1, orange, FD = 0.2, etc. (these are arbitrary cutoffs simply for visualization). These points are then duplicated on the upper bound of the graph so that the various thresholds can be easily compared against the Gray Plots. The idea here is that the proper threshold for removing movement corrupted data should line up with the corrupted data visualized by the Gray Plot (An example is provided in Figure 4). Pre/Pos reg = Pre/Post regression.

In Figure 4 we show a typical “medium mover” participant in the ABCD study. What can be seen in the augmented Gray Plots for Figure 4A is that large spikes in the movement estimates correspond nicely to artifacts in the data that can be visualized in the Gray Plot, and the other quality measurements: DVARS, whole brain signal, and whole brain signal standard deviations (see black arrows, Figure 4A). These are the same head-motion related BOLD signal artifacts that have previously been documented (Power et al., 2014; Power, 2016). However, what one can also visualize in these plots is that if we take the same threshold (FD > 0.2) and examine other places along the FD trace, much of the multiband data that crosses the upper threshold of 0.2, does not appear to have the same types of movement induced artifacts (see red arrows, Figure 4A). These qualitative observations suggest a potential mismatch in movement related artifact in the BOLD data, and movement as reflected by the motion traces specific to the multiband data.

**Figure 4.**
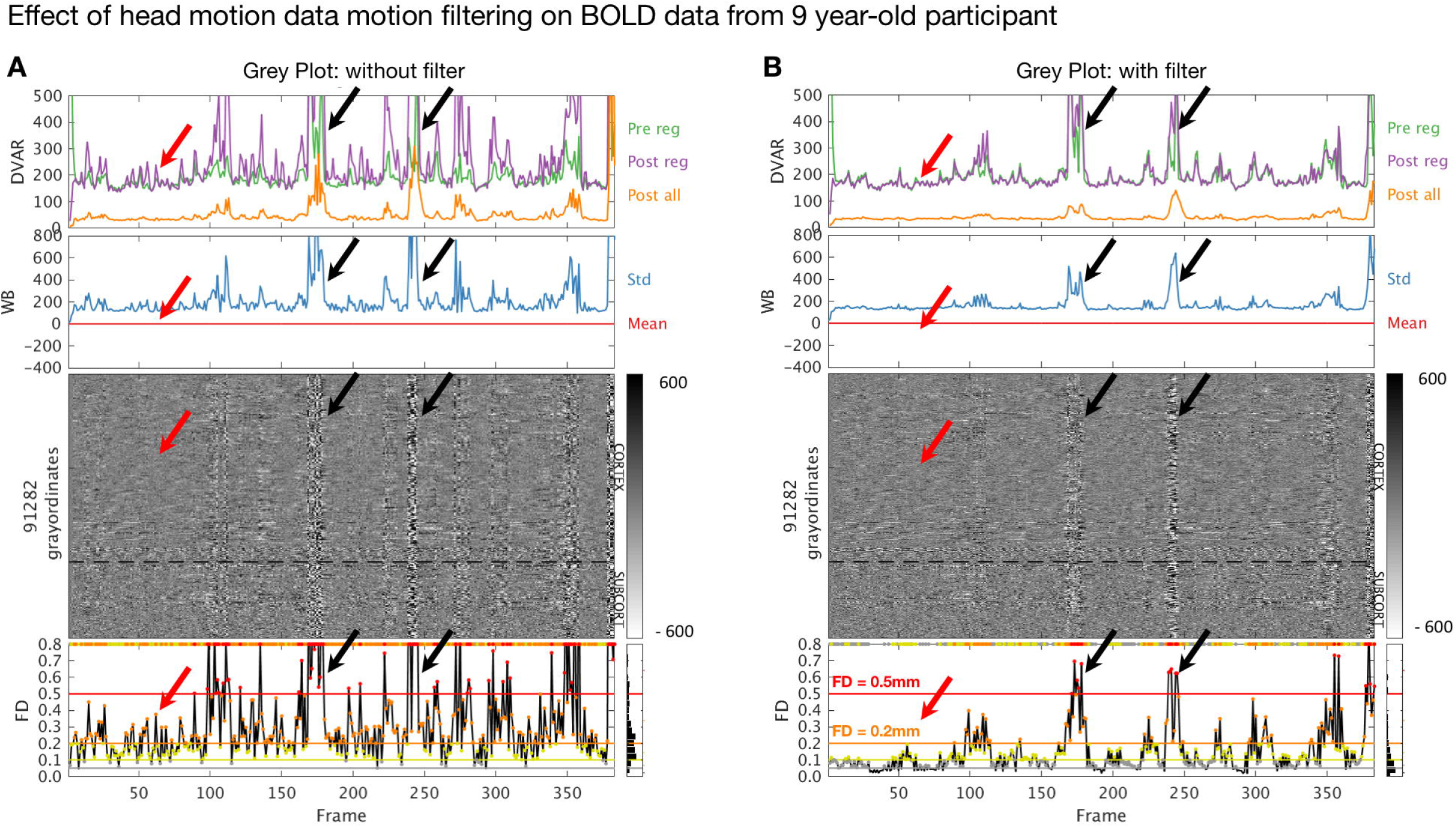
Augmented Gray Plots of a 9-year old ABCD participant. (A) Data through processing where no filter has been applied to the motion estimates. (B) Data through processing where the filter (FF filter) has been applied to the motion estimates. Black arrows highlight large movement spikes that cause artifacts in the BOLD data. After the filter, these spikes remain readily apparent on the motion trace and are associated with artifact. Red arrows highlight smaller spikes of apparent motion without associated artifact in the BOLD data. After the filter, these spikes in the motion data no longer cross commonly used motion thresholds (i.e., FD < 0.2 – brown arrow; and FD < 0.5 – red arrow).

### Multiband imaging reveals previously unrecognized distortions of FD calculations

Having established the possibility of differences in motion estimates for multiband data compared to single band data, we next compared respiratory traces and power spectra of motion estimates in the two data types. Figure 5A illustrates motion parameter timeseries (3 translations, 3 rotations) together with respiratory belt data acquired in a representative ABCD subject. This subject was scanned with the standard ABCD fMRI protocol (MB 6; TR 800ms). Figure 5A (middle row) provides the full trace of a run for a subject, along with a small ‘snap shot’ of 10 frames of that same run (top row). We also provide power spectra for those signals (bottom row). A relation between apparent head motion and the respiratory belt is present but not easily perceived in conventional time series plots. This relation is most clearly evident in the power spectral density plots. Chest wall motion is regularly periodic. Accordingly, the respiratory belt power spectrum (gray trace) shows a well-defined peak (at ∼0.37 Hz in this participant).

**Figure 5.**
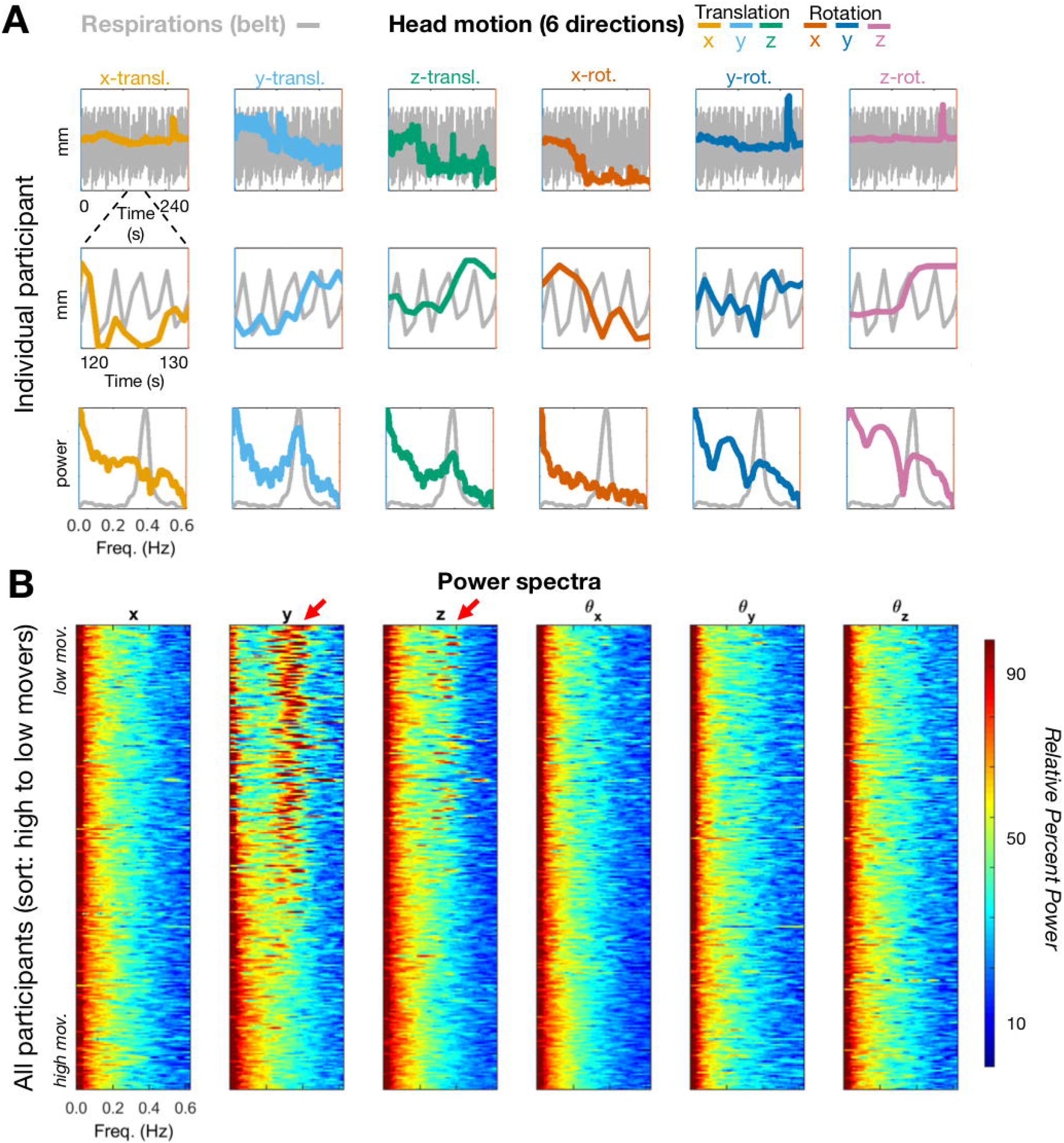
Visualization of motion and respiration power spectra in multiband data. (A) estimates in a single representative ABCD subject (TR = 800ms). Colored lines represent the translation and rotation motion estimates. Top row – full trace of a run for a subject, middle-row – ‘snap shot’ of 10 frames of that same run. Bottom row – power spectrum for both the motion traces and the respiratory trace across runs. (B) power spectra across all participants ranked from lowest to highest movers. Respiration artifact is highlighted with red arrows. To allow for visual comparisons across directions and conditions (i.e. with and without filtering – Figure 5, 6 and SI Figure 2, 3) the color map is a logarithmically transformation to represent the relative percent power of each scan per frequency (see more details in appendix B). The scans are sorted by mean FD.

This peak is mirrored in the y-translation power spectrum and, to a lesser extent, also in the z-translation power spectrum. While the largest correspondence occurs in the y-direction, as would be expected, correspondence in other directions might relate to the fact that the relative head position of the subject often changes from the beginning to the end of a run. It might also represent true head motion related to the respirations themselves.

Figure 5B shows motion parameter power spectra in all participants represented as a stack ordered according to total motion (mean *FD*). The lowest movers are on top. Oscillations at ∼0.3 Hz, predominantly in the y-translation parameter, are evident in the lowest movers. Note the absence of low frequency power particularly in the y-translation column (i.e., no evidence of slow head elevations) in low movers. The ∼0.3 Hz oscillation becomes progressively less prominent towards the bottom of the stack. This effect arises because the power spectra are normalized (equal total power in all traces). Thus, high motion manifests as a shift of relative power towards low frequencies. In other words, factitious y-translation oscillations, if present, are overwhelmed by true high amplitude head motion. Modest uncertainty regarding the direction of true head motion arises from potential cross-talk between the rigid body parameters represented in Eq. (3). These algebraic complications are detailed in Appendix A.

### Respiratory artifact is present, but less prominent in single band data

Figure 6 is identical to Figure 5 except that it shows single-band instead of multiband results. Respiratory belt data were not available in this case. Nevertheless, it can be deduced that respiratory artifact occurs primarily in the y (phase encoding) direction. A peak in the power spectrum is present. However, in comparison to the multiband case, this peak is much broader and centered at ∼0.14 Hz. These power spectrum differences between single and multiband data likely occur because the sampling rate (TR) is not fast enough to capture the true rate of the respirations. Rather, respirations are aliased into other frequencies (see Eq. (5)). This effect is illustrated in SI Figure 1. Any signal that has a frequency higher than half the sampling rate (Nyquist limit) will be detected as a signal of lower frequency. For example, a 2hz signal sampled at 10hz will read out at 2hz. A 5hz signal sampled a 10hz, will read out at 5hz because it is exactly at the Nyquist limit. However, a 6hz signal sampled at 10hz, will not read out as 6hz, but will alias into a lower frequency. In this example, the 6 Hz signal aliases at 4hz.

**Figure 6.**
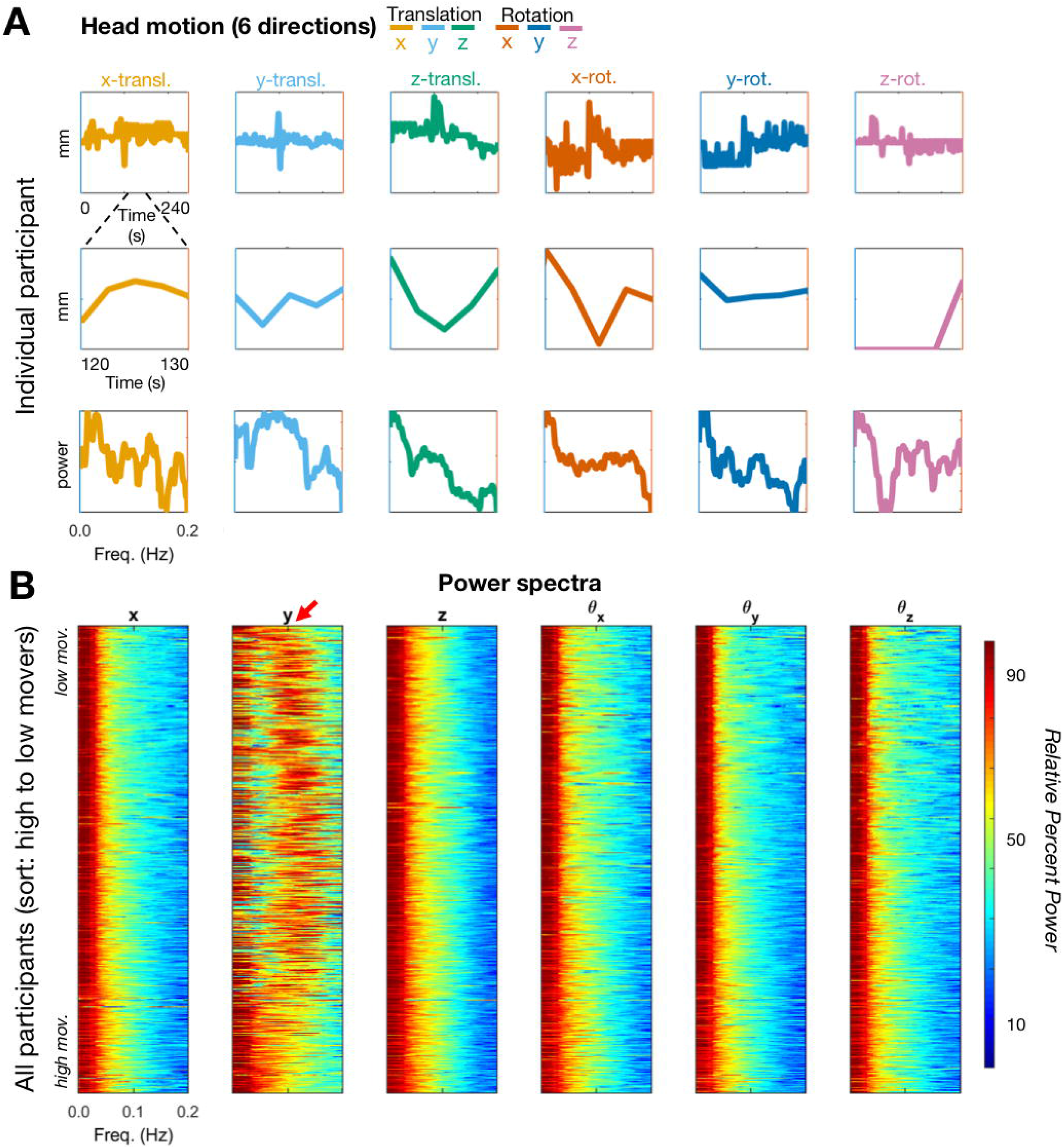
Visualization of motion and respiration power spectrum in single band data. (A) Head motion estimates in a single representative subject (TR = 2.5s). Top row – full trace of a run for a subject, middle-row – ‘snap shot’ of 10 frames of that same run. Bottom row – power spectrum for the motion traces. (B) power spectrum across all participants ranked from lowest to highest movers. To allow for visual comparisons across directions and conditions (i.e., with and without filtering – Figure 5,6 and SI Figure 2,3) the color map is normalized to represents the relative percent power of each scan (per frequency) is represented in dB (log10 transformed and multiplied by 10). The scans are sorted by mean FD.

In SI Figure 1 we provide an illustration of what frequencies might be expected for a given respiratory rate and for a given sampling rate (TR). Three examples are given for 10, 18, and 25 breaths per minute. What is shown is that the artifact in the motion traces for a TR of 800ms (or 0. 8s) will match the true respiratory rate of a given participant; however, for slower TRs (singleband data) the respiratory rate will be aliased into different frequencies. Thus, any variation in respiratory rate during a scan is likely to be spread into different frequencies, broadening the spectral peak (see Figure 6B – red arrow, relative to Figure 5B – red arrow). This mismatch between true respiratory rate and what can be observed in motion traces might be one reason why these artifacts were initially overlooked in single-band data.

An experimental demonstration of the above-described effects is presented in Figure 7. Resting state fMRI was acquired in single subject while respiration was visually paced at exactly 0.33 Hz (see supplementary movie to view the pacing display). The identical multiband sequence was run at TR = 0.8, 1.5, and 2 seconds. fMRI was collected also using a single-shot sequence at TR = 2.5. Aliasing of the respiratory signals in Fig. 7A-D closely matches the theory expressed in Eq. (5) and illustrated in Fig. 7E. Note also that the respiratory peak is narrow because it was precisely paced.

**Figure 7.**
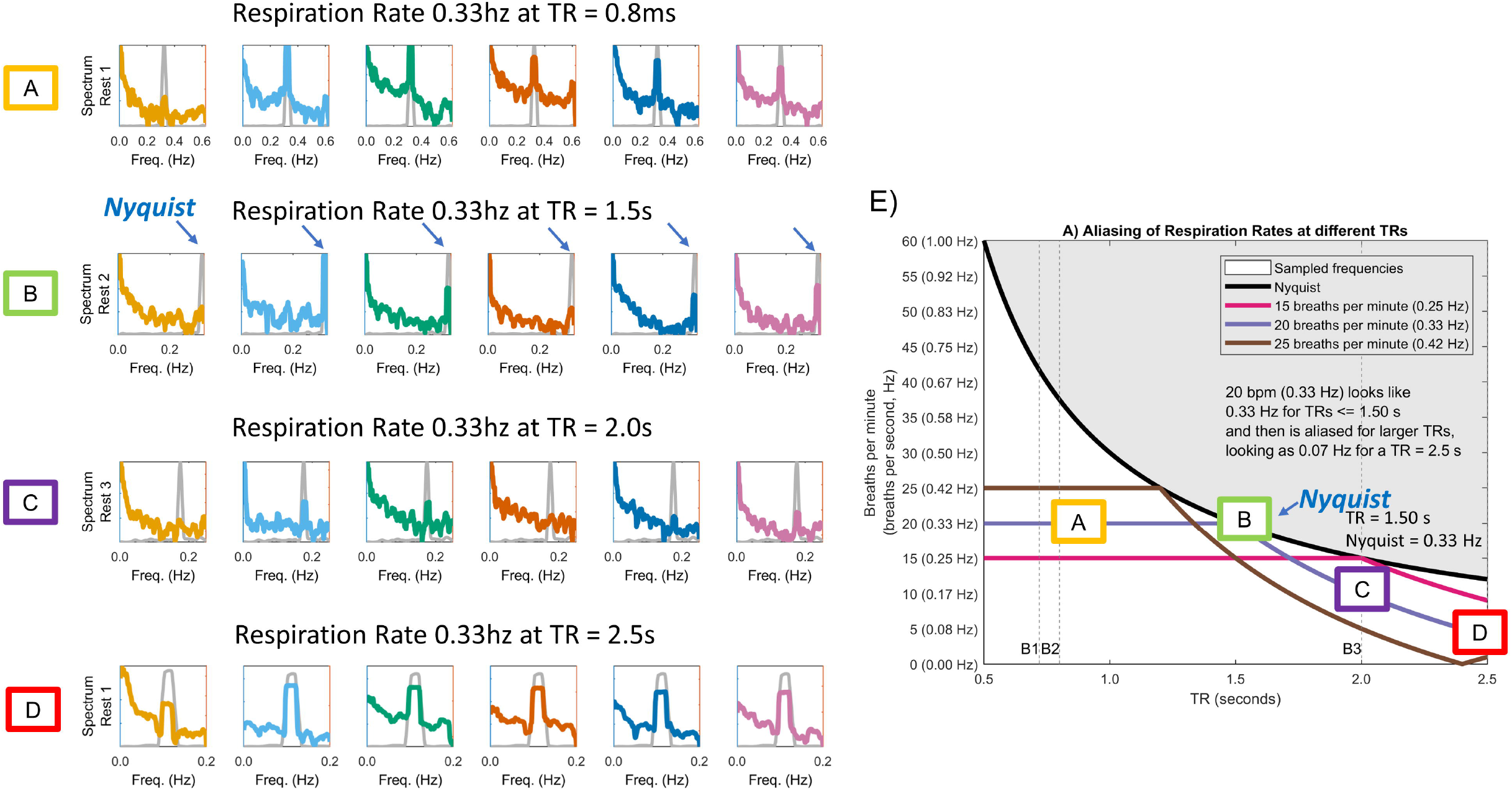
Demonstration of respiration artifact aliasing as a function of sampling rate (TR). BOLD data were collected while the participant’s respiratory rate was visually paced at exactly 0.33hz (see supplementary movie 1). An identical multiband sequence at TR = 1, 1.5, and 2 seconds was used. Data were also collected using single-band sequence at various TRs (A-D) Power spectra for both the motion traces and the respiratory belt trace across runs. (E) Superimposition of empirical spectral peak values over predicted values based on Nyquist theory as outlined in Methods. Note close match of empirical vs. theoretical spectral peak frequencies.

### Filtering FD data corrects for respiratory artifacts and improves estimates of data quality

Having established the effects of respirations on motion estimates, we then turned our attention to lessening this influence. To reduce the influence of factitious head motion, we applied a bandstop/notch filter to the rigid body parameters [Eq. (3)]. Two filter strategies were evaluated.

First, we evaluated a generic, Fixed Frequency [FF] filter designed to exclude a range of respiratory rates (see Methods). The FF approach does not require respiratory rate measurement in individuals. In the second approach, we used respiratory belt data (see Methods) to generate individually tailored filter parameters (Subject Frequency [SF] filter). The effects of applying the FF filter are shown in Figure 4B.

Marked improvement in the linkage between FD and data quality (gray ordinates block) is evident on comparison of Figures 4A vs. 4B (black arrows). The impact of this improvement on FD-based frame censoring is dramatic. Thus, assuming a censoring criterion of > 0.2mm (orange line in FD trace), very few frames would survive without filtering (panel A). In contrast, with filtering (panel B), censored frames align more strongly with the artifact.

Perhaps surprising is that the DVARS trace also improved post filtering. The reason for this improvement is that regression of factitious waveforms introduces spurious variance. Thus, main field perturbations caused by chest wall motion may indirectly corrupt fMRI data via distorted nuisance regressors. This effect is independent of frame censoring. Hence, there are reasons to remove factitious components from motion regressors even if frame censoring is not part of the analysis strategy.

Supplementary Figure 2 and Supplementary Figure 3 provide the replicates of Figure 5 after the FF and SF filters, respectively. What can be visualized in the Figures is that the filter aligns with the artifact induced by respirations. A close look at both Supplementary Figure 2B (FF filter) and Supplementary Figure 3B (SF filter) shows that both these methods do not perfectly capture the artifact. Indeed, in Supplementary Figure 3B, particularly for high moving subjects (i.e., bottom of graph), the filter impinges on true motion values. In other words, the identification of the dominant respiratory rhythm in high movers is difficult. This difficulty represents a barrier to use of the SF strategy in high movers (see Supplementary Figure 4).

### Filtered head motion estimates improve data quality

To quantify the impact of filtering motion parameters on BOLD data quality, we used methodology introduced by Power et al., (2014). The method was originally used to select the frame-censoring criterion using a rational procedure. Here, we adapt the method to assess the fidelity of a quality assurance measure, specifically, FD. As outlined in Figures 1 and 2 (Methods), the procedure involves (1) reordering frames (volumes) according to decreasing quality (greater FD), (2) passing a sliding window through the quality-ordered data (i.e., frames 1-151, 2-152, 3-153, etc.) and calculating the correlation matrix for each window, (3) defining a baseline correlation matrix as the average over the first 30 windows, (4) computing the similarity between the baseline correlation matrix and matrices obtained in subsequent (lower quality) windows. Plotting this similarity vs. [quality ordered] window index reveals the FD value that rationally separates “low” vs. “high” quality data. The statistical significance of the FD criterion is computed against a null distribution generated by applying steps 1-4 to surrogate data in which frame order has been randomized.

Results obtained with the above outlined procedure are shown in Figure 8. The basic result is that filtering causes the boundary between “low” vs. “high” quality data to shift towards lower FD values. In other words, without filtering, relatively high FD values do not reliably indicate compromised data quality. In other words, the unfiltered FD measurements are more random than FD measurements that have been filtered using the FF filter (or SF filter – see Supplementary Figure 4). The top row of Figure 8A demonstrates this effect in a single subject, where fewer green points (windows similar to the baseline) extend up to the highest FD values.

**Figure 8.**
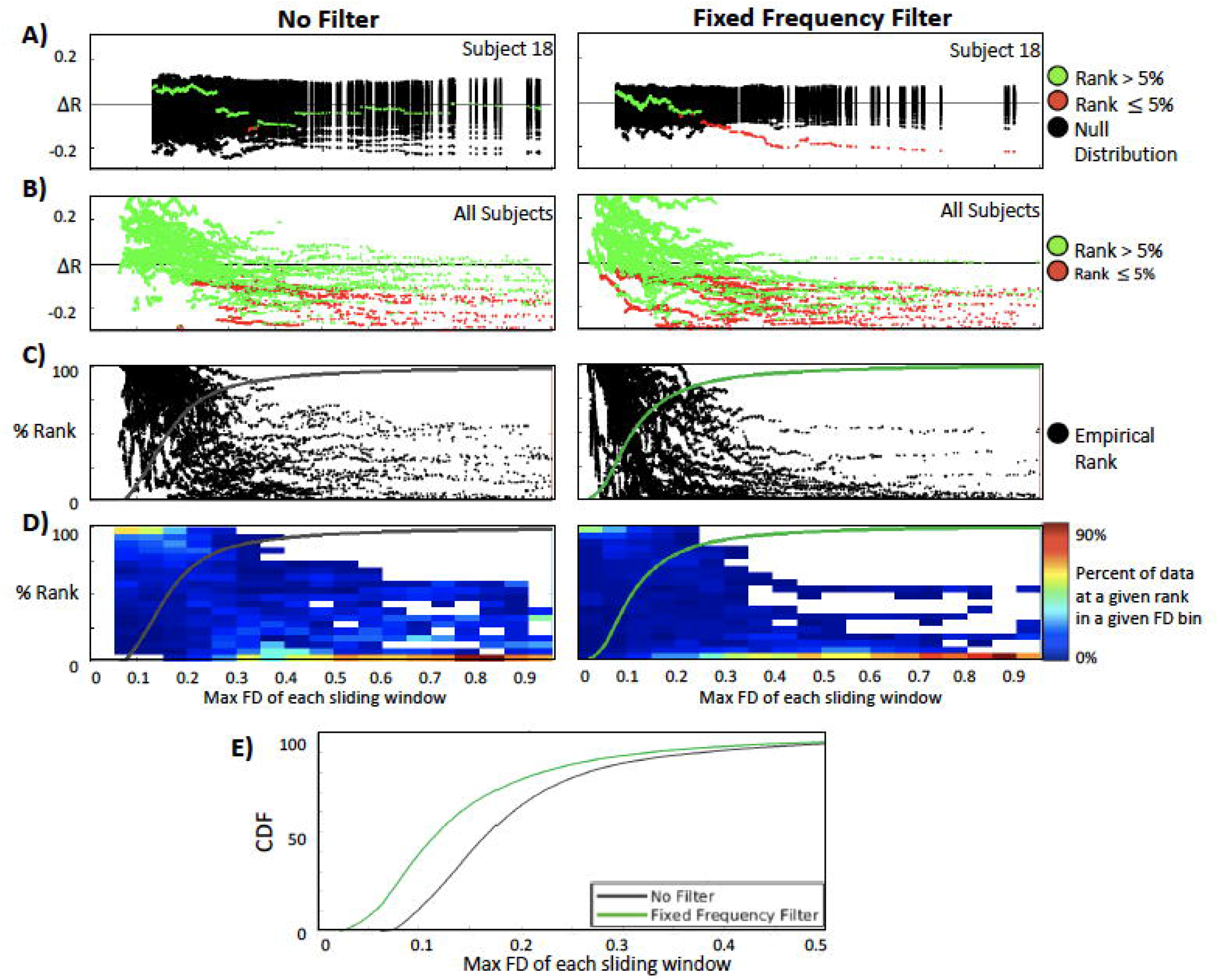
Rank distributions and CDF comparisons of filter conditions. (A) Significance of all FD-value matrices for one subject and for the FF and ‘no filter’ conditions (also see Figure 2). Values in green do not significantly differ from random. Values in red are significantly different from random. For simplicity of visualization, the true and null distributions are zero centered. (B) Significance of all FD-value matrices for all subjects and the FF and ‘no filter’ conditions. Fewer non-significant (green) data points in A and B for the filter conditions shows that the FD estimates are less random. (C) Percentage rank (relative to null) of the QC-ordered outcomes across subjects, which are binned and represented as a heat map (D). (E) The cumulative distribution function (CDF) across the FF and ‘no filter’ FD bins provides an estimate of the average rate at which the windows deviate from the null model. Direct comparisons of the CDFs using the Kolmogorov–Smirnov (KS) test showed the CDF of the ‘FF filter’ to be significantly different from the ‘no filter’ condition (p< 0.0001). See SI Figure 4 for the SF filter condition.

In contrast, with FF filtering (Fig. 8A), a clear boundary between “low” vs. “high” quality data (sharp transition from greed to red points) occurs at 0.26 mm. Figure 8B plots the rank of the FD-ordered results across subjects (as in Figure 2B). Figure 8C bins those ranks (as in Figure 2C). These plots include all subjects and represent a vertical distribution of ranks (relative to randomly ordered volumes) of sliding window comparisons to the baseline. The cumulative distribution function (CDF) across all FD bins (Figure 8C,D) estimates the average rate at which the windows deviate from the null model (as in Figure 2B,C). Filtering causes the CDF to shift leftwards. This leftward shift reflects, at the population level, better correspondence between FD and data quality, as described above in individuals. Figure 8E plots the CDFs obtained in the FF filter versus the ‘no filter’ conditions. The FF filter CDF significantly differed from the ‘no filter’ CDF (p< 0.0001) by the Kolmogorov–Smirnov (K-S) test. Similar CDF findings were obtained with the SF filter (see Supplementary Figure 4).

### Real-time implementation of head motion parameter filtering

As noted in the Introduction, real-time motion monitoring tools such as FIRMM potentially reduce costs and improve data quality. Here we demonstrate this principle. Real-time filtering of respiratory frequencies has been built into version 3 of FIRMM software. Scanner operators have the option to apply the FF filter in real time (Figure 9 and SI Movie 2) via a GUI that provides users with control over filter parameters. This type of control is necessary as different populations (e.g., children vs. adults) have different respiratory rates. Figure 9 shows the FIRMM display for unfiltered and filtered data acquired in three 5-minute resting state BOLD fMRI runs. As in our original report (Dosenbach et al, 2017), on-line filtered FD values very closely track filtered FD values generated by the off-line processing stream (Figure 9C,D).

**Figure 9.**
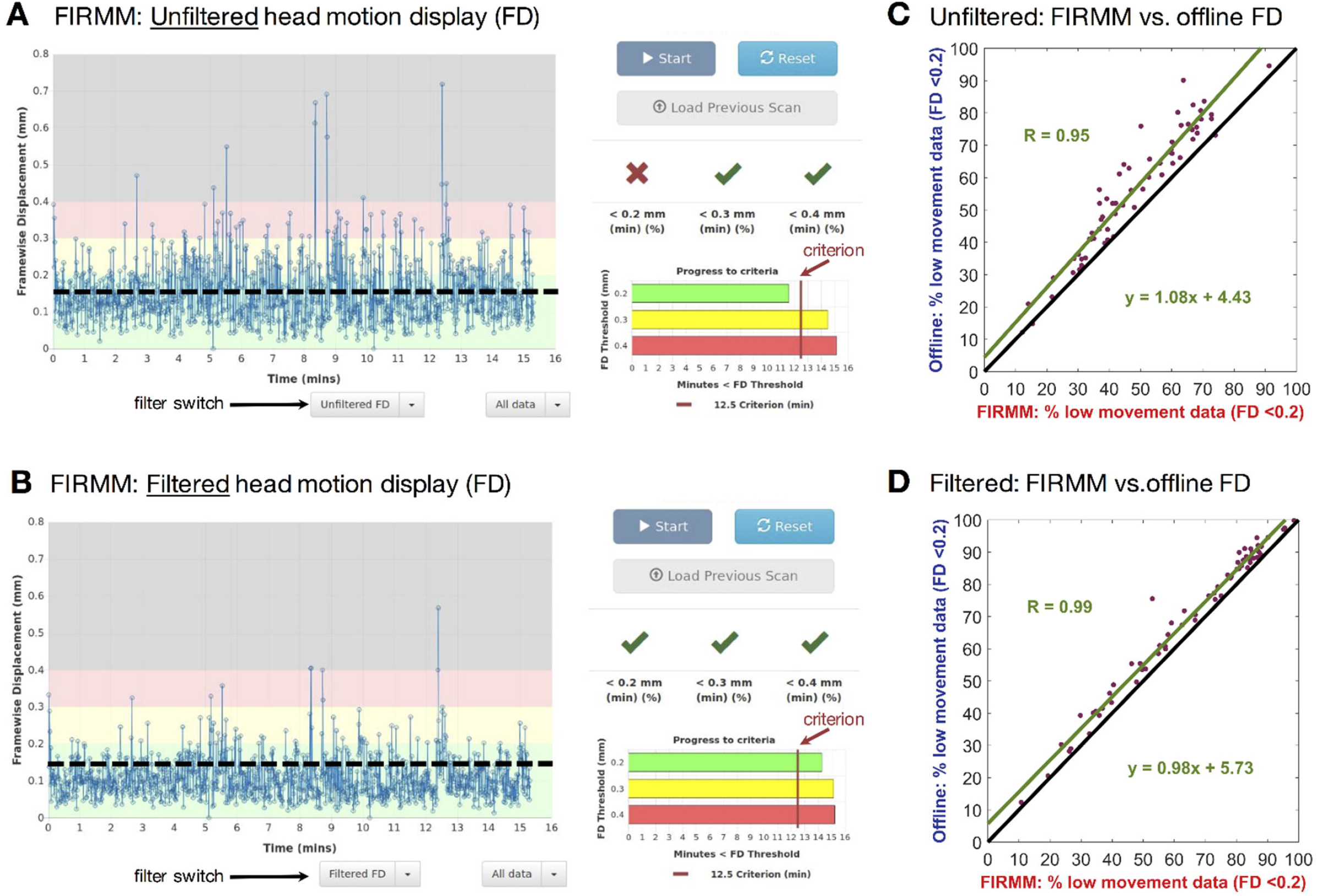
FIRMM display of unfiltered (A) and filtered data (B) for a subject with 3 × 5-minute resting runs. (C and D) Comparisons of percentage of usable frames from FIRMM compared to post-processing utilized here (mcflirt) showed highly significant correspondence. FIRMM was conservative relative to offline post-processing in that it was more sensitive to ‘bad’ frames by approximately 5% of the total.

## Discussion

We show that respiratory motion artifactually inflates head motion estimates. A consequence is overly aggressive frame-censoring and unnecessary data loss. Faulty head motion estimation also interferes with other artifact suppression techniques apart from frame-censoring. Specifically, corruption of nuisance regressors derived from retrospective head motion correction introduces spurious variance into the fMRI time series. This spurious variance appears in DVARS time series and is reduced by appropriate head motion filtering (Fig. 4). Similar logic underlies the rationale for adjusting the spectral content of head motion time series prior to their use as nuisance regressors (Carp, 2011; Hallquist et al., 2013). We demonstrate these principles using on-line as well as off-line head motion estimation based on registration of reconstructed images. However, the same issue potentially arises with any motion estimation strategy. For example, volumetric navigator sequences for structural MRI, which adjust acquisition parameters based on head motion estimates between whole-brain EPI snapshots, might overcorrect during low-movement scans (Zaitsev et al., 2017). To address these important concerns, we evaluated a notch filter that drastically reduces the effects of respirations on FD estimates and thus improves MRI quality. Finally, we demonstrate near real time implementation of on-line motion estimation including notch filter correction of FD estimates. This development represents an important improvement of our existing real-time motion monitoring software.

### Distribution of respiratory motion across rigid body parameters

Although respiratory head motion, likely factitious in origin, prominently occurs in the phase encoding direction (Figs. 5,6), head motion at the same frequency may appear also in any motion parameter (e.g., Fig. 7D). Respiratory head motion affecting parameters other than phase encoding direction displacement can arise on the basis two distinct mechanisms that must be distinguished in future work. First, true respiratory motion can be transmitted from the chest to the head by mechanical linkage through the neck, as discussed above in connection with Fig. 5. This motion is most likely to occur as z-direction displacement (as in Figs. 5 and 6) or rotation about the ear-to-ear axis (*θ_x_*; pitch). Being true, this motion may generate BOLD signal artifact on the basis of spin history effects (Friston et al., 1996a). Second, factitious head motion may appear in any motion parameters owing to algebraic cross-talk (see Appendix A for details). In this case, the artifact would appear to bleed into the other directions. It is for this reason we chose to apply our notch filter to all directions of head motion. Nonetheless, respirations contaminate BOLD data in multiple ways (Birn et al., 2006; Power et al., 2017); Additional work will be needed to disentangle these complex issues.

### Pros and cons of the SF and FF filters

We empirically designed the FF notch filter to capture a large proportion of respiratory frequencies in the ABCD participant sample (see Methods). This filter worked well in improving the current functional connectivity results. However, as different populations and age groups have different respiratory rates (Wallis et al., 2005), the FF filter should be tailored to the population being studied. Accordingly, the FIRMM software enables the user to modify the FF filter parameters via the GUI. Also, we note that some populations might not require motion parameter filtering at all. In particular, infant chest excursions are likely to only minimal perturb the main field. For this reason, FIRMM allows the user to control whether filtering is enabled or disabled.

In principle, the SF filter strategy, which matches the filter characteristics to each individual’s respiratory rate, should, more specifically than the FF filter, eliminate factitious components from the head motion regressors. In the current results, the SF filter performed similarly to the FF filter in low movers. In high movers, the SF filter strategy often failed to correctly identify the respiratory component (Supplementary Figures S3 and S4). Thus, while the SF filter is appealing, there are potential pitfalls. First, not all studies collect respiratory belt data, which is required for using a SF filter. Second, even if respiratory data were collected on a subject, they could be inaccurate. Often the respiratory belts are not fit properly or become dislodged during scanning. Along the same lines, identifying the peak respirations from the motion numbers themselves can also be problematic, as noted above. Future work is needed to develop reliable SF filtering. One option might simply be to modify the notch filter applied here to a low-pass filter where identifying the proper range of threshold would not be required (Siegel et al., 2016).

Importantly, this manuscript did not consider the potential need for adjusting thresholding standards for frame censoring. Applying either the FF or SF filter necessarily reduces overall power of the FD traces. This reduction in power likely requires a revisiting of the thresholds for ‘scrubbing’ that have become common practice in the literature (e.g., FD < 0.2) (Power et al., 2014). While beyond the scope of the current report, a systematic evaluation to this end is needed in future work.

### FIRMM allows scanner operators to choose filter parameters based on the population being studied

FIRMM was initially created as a response to difficulties in obtaining enough quality ‘movement free’ data collection for t-fMRI and rs-fcMRI studies. FIRMM calculates and displays curated motion values (i.e., frame displacement [FD]) and summary statistics during data acquisition in real time, providing investigators and operators estimates of low-movement data. The benefits of using FIRMM to reduce cost savings and improve data quality have been thoroughly examined in prior reports (Dosenbach et al., 2017; Greene et al., 2018). Importantly, FIRMM and other real-time motion estimators rely on accurate translation and rotation numbers. The implementation of the notch filter in real time (see methods for our approach) improves the accuracy of the motion values in FIRMM.

There are a few considerations for users. As noted in the methods, when applying our FIRMM notch filter to the motion estimates. The filter recursively weights the two previous samples to provide an instantaneous filtered signal, and thus cannot start until the third sample. In addition, to avoid a phase delay with respect to the original signal, the filter is applied again in the reverse direction (cancelling the phase delay). In real time, this requires waiting until the 5^th^ sample to provide a best estimate of motion post filter for the third frame. Each time a new frame arrives the process is reapplied to all frames available, continually converging closer to the optimal output obtained when the filter is applied twice to the entire sequence (as in post-processing). This procedure works well and only leads to a small delay in motion estimates of at least two frames. For a large majority of users this small delay for fast TRs (< 1 sec/frame) accompanying multiband imaging will be non-consequential. Nevertheless, alternative real-time filtering algorithms exist that theoretically could reduce or eliminate this delay (Glover Jr, 1977). Implementation of such filters will be considered in the future.

### Respiratory artifacts in motion estimates are not isolated to SMS sequences

Although it has been known for over a decade that respiratory motion perturbs the main field (Van de Moortele et al., 2002) these effects have gone largely ignored. Nevertheless, these effects materially compromise denoising strategies, especially with SMS sequences (e.g., Fig. 4). As fast fMRI with SMS sequences becomes the standard of practice, large, ongoing, multisite projects, such as Adolescent Brain Cognitive Development (ABCD), must consider this issue. Here, we demonstrate one approach to managing this problem.

Similar corruption of head motion estimates in single-band data has not been widely recognized. Reasons for this circumstance include that aliasing (Figures 7 and S1) disguises the source of the problem and that respiratory rate variation normally spans a comparatively large fraction of the resolvable spectral band (Figure 6). However, factitious respiratory motion artifact can be detected in single-band data. Although we did not systematically examine the effects of applying our notch filter to single band data, doing so might improve previously obtained results. We note that this may not be straightforward as aliasing [(Eq (5)] must be taken into account. As noted above, one option, might be to simply apply a low-pass filter, instead of a band-stop or notch filter (Gordon et al., 2017; Gratton et al., 2018; Laumann et al., 2016).

## Conclusions

Properly controlling for motion artifacts in fMRI is of the utmost importance in both clinical and research applications. The current report shows that main field perturbations generated by chest wall motion corrupt measurement of head motion. Factitious head motion leads to the false appearance of compromised data quality and introduces spurious variance into fMRI time series via corrupted nuisance regressors. These effects are readily apparent in SMS sequences but are also present in single-band sequences. Notch filtering tailored to the respiratory frequency band improves the accuracy of head motion estimates and improves functional connectivity analyses. Notch filtering can be applied off-line during post-processing as well as on-line to improve real-time motion monitoring. This approach might also have merit to improve recently introduced volumetric navigator motion correction procedures for structural T1 and T2 scans as well (Tisdall et al., 2016), as these techniques utilize within run EPI snapshots to monitor head position. Additional work is needed to improve strategies for tuning filters to better match individual respiratory rates.

## Acknowledgements

We would like to thank the ABCD MRI image acquisition working group, including BJ Casey and Jonathan Polimeni, for their work in organizing the group and developing the ABCD MRI acquisition protocols, respectively. We would also like to thank the other massive contributions to the ABCD effort by the other PIs and staff (see https://abcdstudy.org/). This work was supported by the National Institutes of Health (grants R01 MH096773 and K99/R00 MH091238 to D.A.F., R01 MH115357 to D.A.F, J.T.N., R01 MH086654, J.T.N., U24 DA04112 to A.D., U01 DA041148 to D.A.F., S.W.F., B.J.N), the Oregon Clinical and Translational Research Institute (D.A.F), the Gates Foundation (D.A.F), the Destafano Innovation Fund (D.A.F.), a OHSU Fellowship for Diversity and Inclusion in Research Program (O.M.-D.), a Tartar Trust Award (O.M.-D.), the OHSU Parkinson Center Pilot Grant Program (O.M.-D.) and a National Library of Medicine Postdoctoral Fellowship (E.F.).

---

**Supplementary Figure 1.**
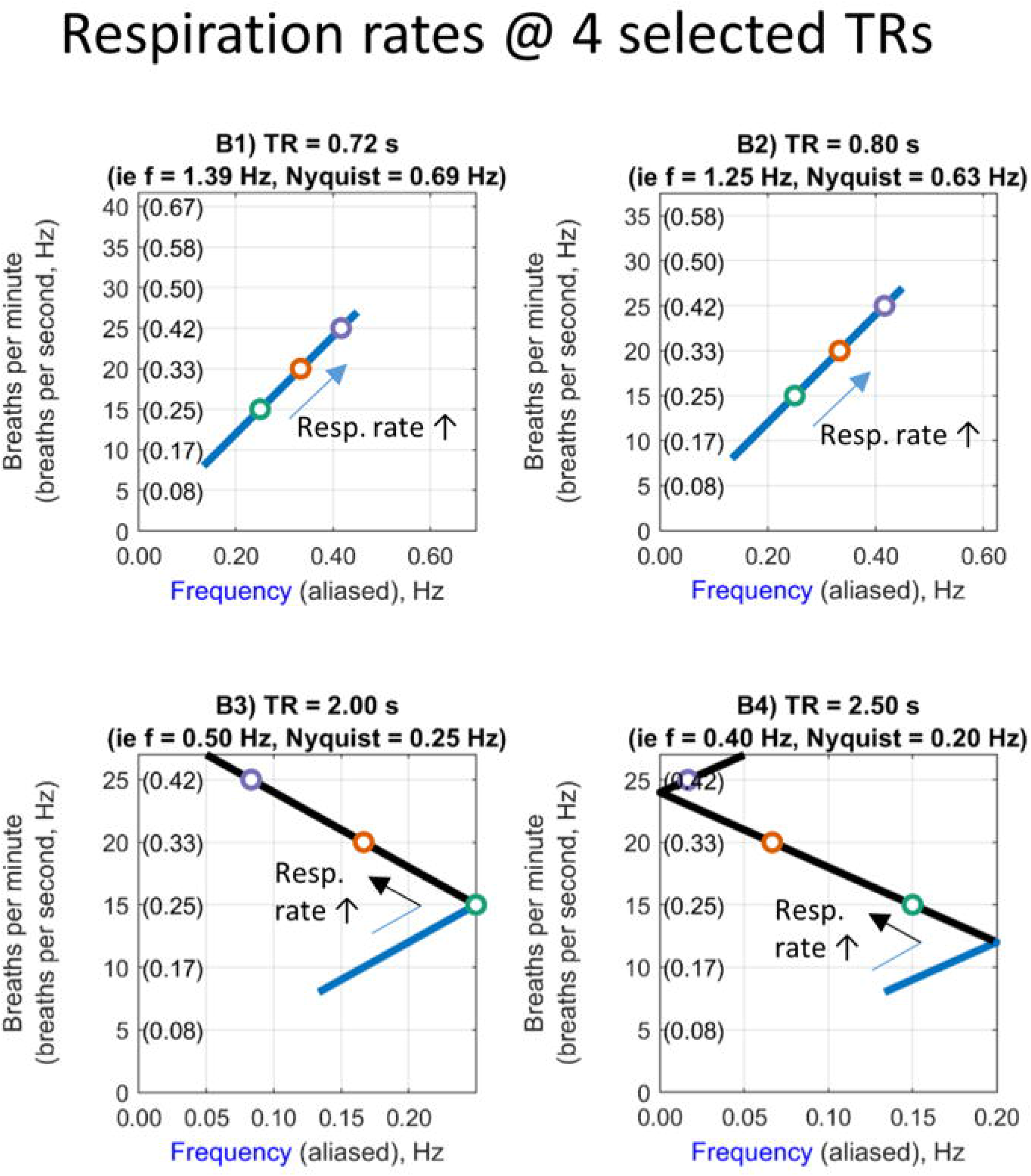
Aliasing due to the slow sampling rate of single band data, relative to respiration rate at 4 selected TRs.

**Supplementary Figure 2.**
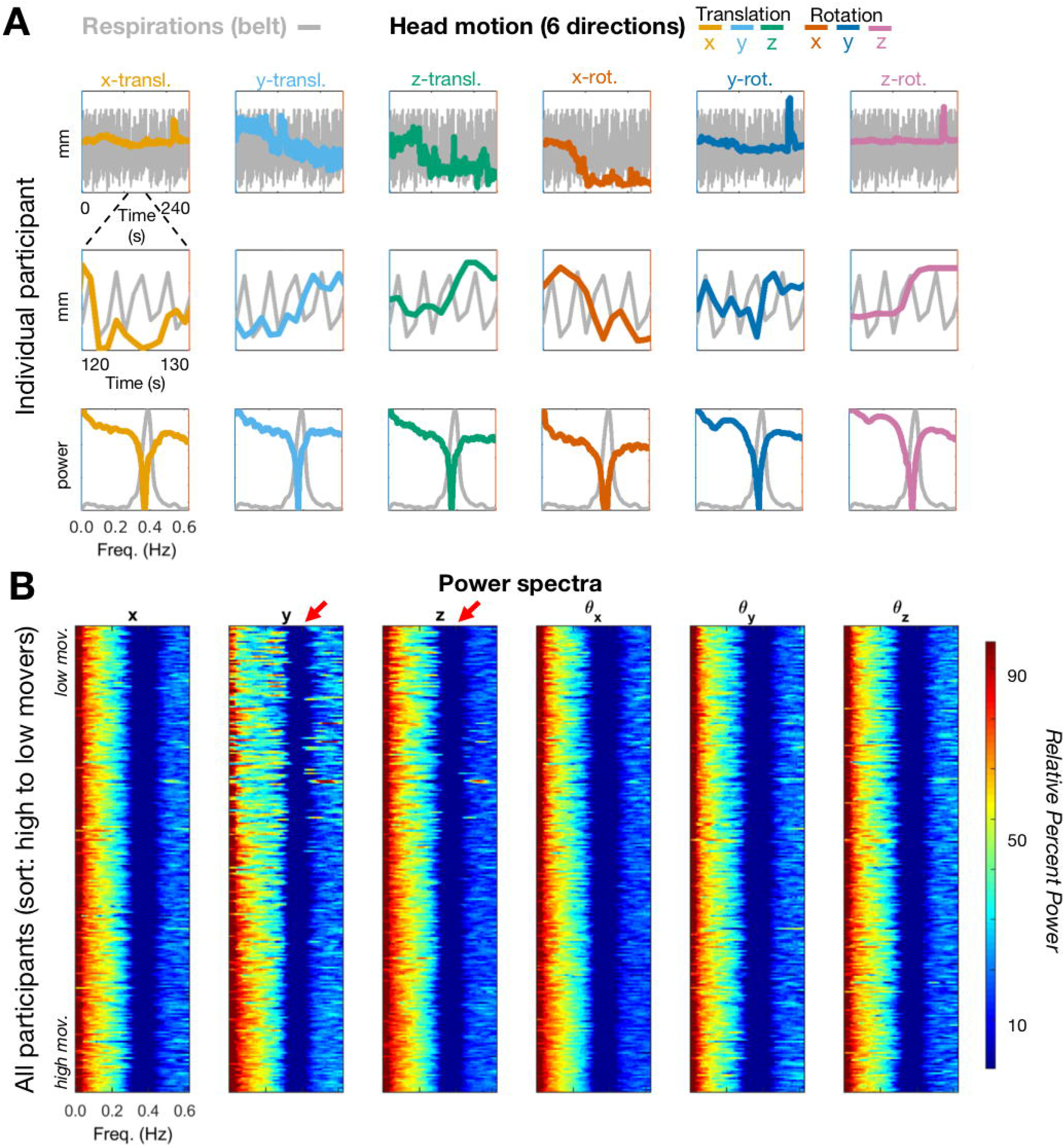
Visualization of motion and respiration power spectrum in multiband data after the FF filter. (A) estimates in a single representative ABCD subject (TR = 800ms). Top row – full trace of a run for a subject, middle-row – ‘snap shot’ of 10 frames of that same run. Bottom row – power spectrum after filter for both the motion traces and the respiratory trace across runs. (B) Filtered power spectra across all participants ranked from lowest to highest movers. The result of filtering on the respiration artifact is highlighted with red arrows. To allow for visual comparisons across directions and conditions (i.e. with and without filtering – Figure 5,6 and SI Figure 2,3) the stacked spectra are represented in dB as in Fig. 6. The subjects are sorted by mean FD (lowest movers on top).

**Supplementary Figure 3.**
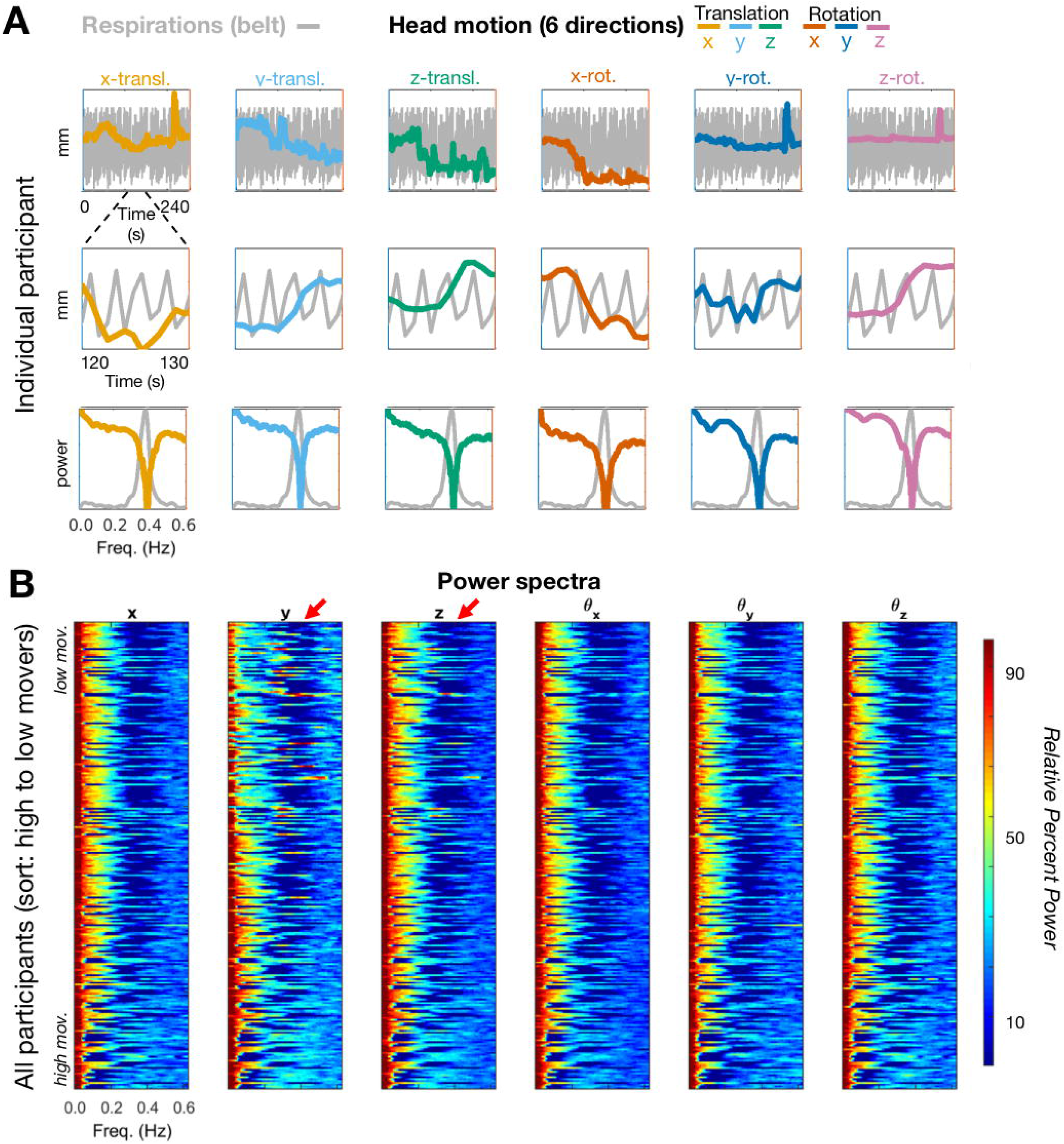
Visualization of motion and respiration power spectra in multiband data after SF filter. (A) Same ABCD participant illustrated in Fig. S2 (TR = 800ms). Top row – full trace of a run for a subject, middle-row – ‘snap shot’ of 10 frames of that same run. Bottom row – power spectrum after filter for both the motion traces and the respiratory trace across runs. (B) Filtered power spectra across all participants ranked from lowest to highest movers. Respiration artifact is highlighted with red arrows. Conventions as in Fig. S2.

**Supplementary Figure 4.**
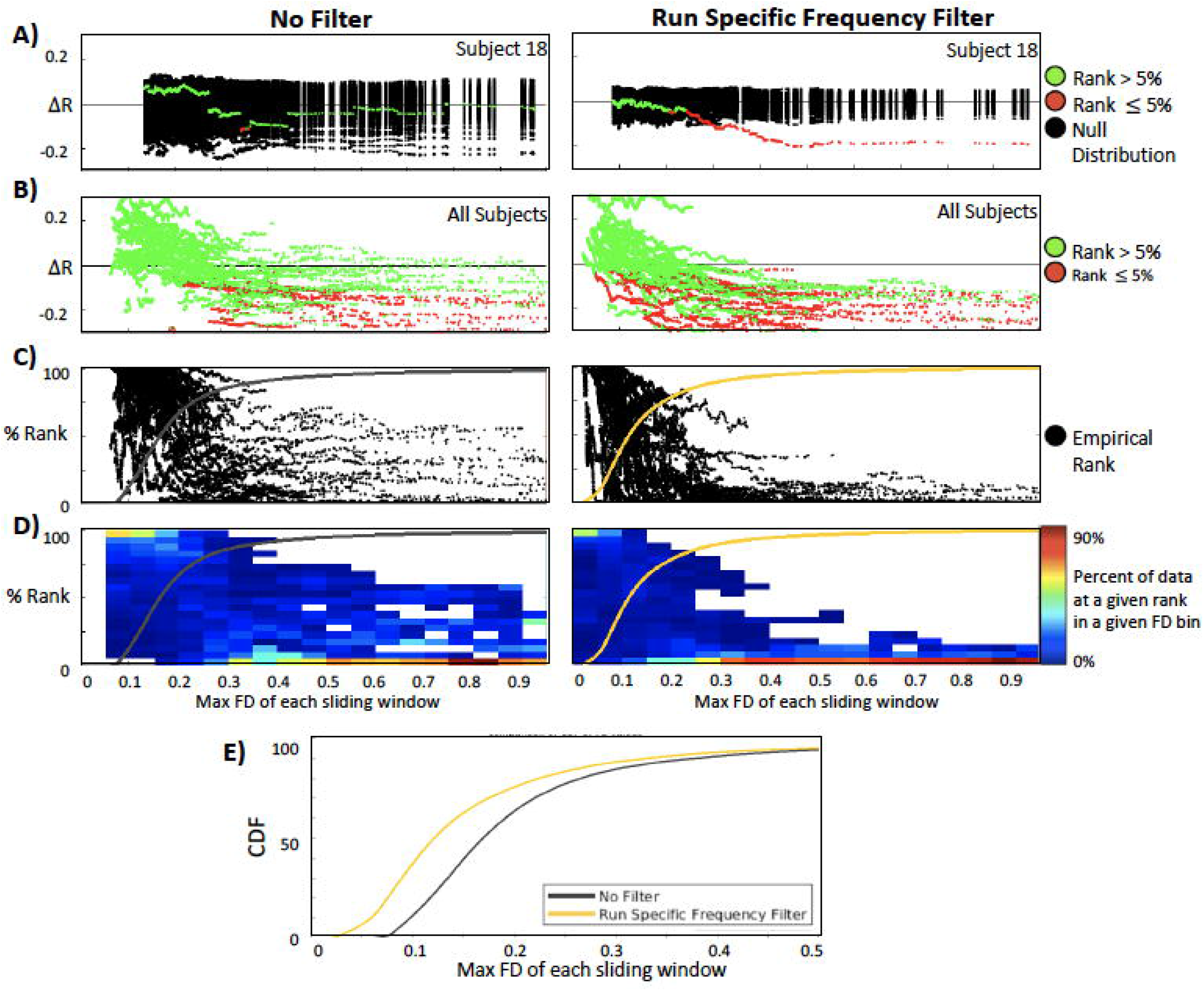
Rank distributions and CDF comparisons of filter conditions. (A) Significance of all FD-value matrices for one subject and for the SF and ‘no filter’ conditions (also see Figure 2). Values in green do not significantly differ from random. Values in red are significantly different from random. For simplicity of the visualization, the true and null distributions are zero centered. (B) Significance of all FD-value matrices for all subjects and the SF and ‘no filter’ conditions. Fewer non-significant (green) data points in A and B for the filter conditions shows that the FD estimates are less random. (C) Percentage rank (relative to null) of the QC-ordered outcomes across subjects, which are binned and represented as a heat map (D). (E) The cumulative distribution function (CDF) across the SF and ‘no filter’ FD bins provides an estimate of the average rate at which the windows deviate from the null model. Direct comparisons of the CDFs using the Kolmogorov-Smirnov (KS) test showed the CDF of the ‘SF filter’ to be significantly different from the ‘no filter’ condition (p< 0.0001). See Figure 8 for the FF filter condition.

## References

Barth, M., Breuer, F., Koopmans, P.J., Norris, D.G., Poser, B.A., 2016. Simultaneous multislice (SMS) imaging techniques. Magn. Reson. Med. https://doi.org/10.1002/mrm.25897

Behzadi, Y., Restom, K., Liau, J., Liu, T.T., 2007. A component based noise correction method (CompCor) for BOLD and perfusion based fMRI. Neuroimage 37, 90–101. https://doi.org/10.1016/j.neuroimage.2007.04.042

Birn, R.M., Diamond, J.B., Smith, M.A., Bandettini, P.A., 2006. Separating respiratory-variation-related fluctuations from neuronal-activity-related fluctuations in fMRI. Neuroimage.

Bjork, J.M., Straub, L.K., Provost, R.G., Neale, M.C., 2017. The ABCD Study of Neurodevelopment: Identifying Neurocircuit Targets for Prevention and Treatment of Adolescent Substance Abuse. Curr. Treat. Options Psychiatry 4, 196–209. https://doi.org/10.1007/s40501-017-0108-y

Burgess, G.C., Kandala, S., Nolan, D., Laumann, T.O., Power, J., Adeyemo, B., Harms, M.P., Petersen, S.E., Barch, D.M., 2016. Evaluation of Denoising Strategies To Address Motion-Correlated Artifact in Resting State fMRI Data from the Human Connectome Project. Brain Connect. brain.2016.0435. https://doi.org/10.1089/brain.2016.0435

Carp, J., 2011. Optimizing the order of operations for movement scrubbing: Comment on Power et al. Neuroimage 76, 436–8. https://doi.org/10.1016/j.neuroimage.2011.12.061

Casey, B.J., Cannonier, T., Conley, M.I., Cohen, A.O., Barch, D.M., Heitzeg, M.M., Soules, M.E., Teslovich, T., Dellarco, D. V., Garavan, H., Orr, C.A., Wager, T.D., Banich, M.T., Speer, N.K., Sutherland, M.T., Riedel, M.C., Dick, A.S., Bjork, J.M., Thomas, K.M., Chaarani, B., Mejia, M.H., Hagler, D.J., Daniela Cornejo, M., Sicat, C.S., Harms, M.P., Dosenbach, N.U.F., Rosenberg, M., Earl, E., Bartsch, H., Watts, R., Polimeni, J.R., Kuperman, J.M., Fair, D.A., Dale, A.M., 2018. The Adolescent Brain Cognitive Development (ABCD) study: Imaging acquisition across 21 sites. Dev. Cogn. Neurosci. 0–1. https://doi.org/10.1016/j.dcn.2018.03.001

Chen, L., Vu, A.T., Xu, J., Moeller, S., Ugurbil, K., Yacoub, E., Feinberg, D.A., 2015. Evaluation of highly accelerated simultaneous multi-slice EPI for fMRI. Neuroimage 104, 452–459. https://doi.org/10.1016/j.neuroimage.2014.10.027

Ciric, R., Wolf, D.H., Power, J.D., Roalf, D.R., Baum, G.L., Ruparel, K., Shinohara, R.T., Elliott, M.A., Eickhoff, S.B., Davatzikos, C., Gur, R.C., Gur, R.E., Bassett, D.S., Satterthwaite, T.D., 2017. Benchmarking of participant-level confound regression strategies for the control of motion artifact in studies of functional connectivity. Neuroimage 154, 174–187. https://doi.org/10.1016/j.neuroimage.2017.03.020

Costa Dias, T.G., Iyer, S.P., Carpenter, S.D., Cary, R.P., Wilson, V.B., Mitchell, S.H., Nigg, J.T., Fair, D.A., 2015. Characterizing heterogeneity in children with and without ADHD based on reward system connectivity. Dev. Cogn. Neurosci. 11, 155–174. https://doi.org/10.1016/j.dcn.2014.12.005

Dale, A.M., Fischl, B., Sereno, M.I., 1999. Cortical surface-based analysis. I. Segmentation and surface reconstruction. Neuroimage 9, 179–194. https://doi.org/10.1006/nimg.1998.0395

Di Martino, A., Fair, D.A., Kelly, C., Satterthwaite, T.D., Castellanos, F.X., Thomason, M.E., Craddock, R.C., Luna, B., Leventhal, B.L., Zuo, X.-N., Milham, M.P., 2014. Unraveling the Miswired Connectome: A Developmental Perspective. Neuron 83, 1335–1353. https://doi.org/10.1016/j.neuron.2014.08.050

Dosenbach, N.U.F., Koller, J.M., Earl, E.A., Miranda-Dominguez, O., Klein, R.L., Van, A.N., Snyder, A.Z., Nagel, B.J., Nigg, J.T., Nguyen, A.L., Wesevich, V., Greene, D.J., Fair, D.A., 2017. Real-time motion analytics during brain MRI improve data quality and reduce costs. Neuroimage 161, 80–93. https://doi.org/10.1016/j.neuroimage.2017.08.025

Fair, D.A., Nigg, J.T., Iyer, S., Bathula, D., Mills, K.L., Dosenbach, N.U.F., Schlaggar, B.L., Mennes, M., Gutman, D., Bangaru, S., Buitelaar, J.K., Dickstein, D.P., Di Martino, A., Kennedy, D.N., Kelly, C., Luna, B., Schweitzer, J.B., Velanova, K., Wang, Y.-F., Mostofsky, S., Castellanos, F.X., Milham, M.P., 2012. Distinct neural signatures detected for ADHD subtypes after controlling for micro-movements in resting state functional connectivity MRI data. Front. Syst. Neurosci. 6, 80. https://doi.org/10.3389/fnsys.2012.00080

Fair, D.A., Posner, J., Nagel, B.J., Bathula, D., Dias, T.G.C., Mills, K.L., Blythe, M.S., Giwa, A., Schmitt, C.F., Nigg, J.T., 2010. Atypical default network connectivity in youth with attention-deficit/hyperactivity disorder. Biol. Psychiatry 68, 1084–1091. https://doi.org/10.1016/j.biopsych.2010.07.003

Feczko, E., Balba, N., Miranda-Dominguez, O., Cordova, M., Karalunas, S.L., Irwin, L., Demeter, D.V., Hill, A.P., Langhorst, B.H., Grieser Painter, J., Van Santen, J., Fombonne, E.J., Nigg, J.L., Fair, D.A., 2017. Subtyping cognitive profiles in Autism Spectrum Disorder using a random forest algorithm. Neuroimage. https://doi.org/10.1016/j.neuroimage.2017.12.044

Feinberg, D.A., Yacoub, E., 2012. The rapid development of high speed, resolution and precision in fMRI. Neuroimage. https://doi.org/10.1016/j.neuroimage.2012.01.049

Fischl, B., 2012. FreeSurfer. Neuroimage 62, 774–781. https://doi.org/10.1016/j.neuroimage.2012.01.021

Fischl, B., Dale, A.M., 2000. Measuring the thickness of the human cerebral cortex from magnetic resonance images. Proc Natl Acad Sci U S A 97, 11050–11055. https://doi.org/10.1073/pnas.200033797

Fonov, V., Evans, A.C., Botteron, K., Almli, C.R., McKinstry, R.C., Collins, D.L., 2011. Unbiased average age-appropriate atlases for pediatric studies. Neuroimage 54, 313–327. https://doi.org/10.1016/j.neuroimage.2010.07.033

Friston, K.J., Williams, S., Howard, R., Frackowiak, R.S., Turner, R., 1996a. Movement-related effects in fMRI time-series. Magn. Reson. Med. 35, 346–355.

Friston, K.J., Williams, S., Howard, R., Frackowiak, R.S., Turner, R., 1996b. Movement-related effects in fMRI time-series. Magn. Reson. Med. 35, 346–355.

Gates, K.M., Molenaar, P.C.M., Iyer, S.P., Nigg, J.T., Fair, D.A., 2014. Organizing heterogeneous samples using community detection of GIMME-derived resting state functional networks. PLoS One 9, e91322. https://doi.org/10.1371/journal.pone.0091322

Glasser, M.F., Sotiropoulos, S.N., Wilson, J.A., Coalson, T.S., Fischl, B., Andersson, J.L., Xu, J., Jbabdi, S., Webster, M., Polimeni, J.R., Van Essen, D.C., Jenkinson, M., 2013. The minimal preprocessing pipelines for the Human Connectome Project. Neuroimage 80, 105–24. https://doi.org/10.1016/j.neuroimage.2013.04.127

Glover Jr, J., 1977. Adaptive noise canceling applied to sinusoidal interferences. Acoust. Speech Signal Process. IEEE Trans. 25, 484–491.

Gordon, E.M., Laumann, T.O., Gilmore, A.W., Newbold, D.J., Greene, D.J., Berg, J.J., Ortega, M., Hoyt-Drazen, C., Gratton, C., Sun, H., Hampton, J.M., Coalson, R.S., Nguyen, A.L., McDermott, K.B., Shimony, J.S., Snyder, A.Z., Schlaggar, B.L., Petersen, S.E., Nelson, S.M., Dosenbach, N.U.F., 2017. Precision Functional Mapping of Individual Human Brains. Neuron 95, 791–807.e7. https://doi.org/10.1016/j.neuron.2017.07.011

Gratton, C., Laumann, T.O., Nielsen, A.N., Greene, D.J., Gordon, E.M., Gilmore, A.W., Nelson, S.M., Coalson, R.S., Snyder, A.Z., Schlaggar, B.L., Dosenbach, N.U.F., Petersen, S.E., 2018. Functional Brain Networks Are Dominated by Stable Group and Individual Factors, Not Cognitive or Daily Variation. Neuron 439–452. https://doi.org/10.1016/j.neuron.2018.03.035

Grayson, D.S., Ray, S., Carpenter, S., Iyer, S., Dias, T.G.C., Stevens, C., Nigg, J.T., Fair, D.A., 2014. Structural and functional rich club organization of the brain in children and adults. PLoS One 9, e88297. https://doi.org/10.1371/journal.pone.0088297

Greene, D.J., Koller, J.M., Hampton, J.M., Wesevich, V., Van, A.N., Nguyen, A.L., Hoyt, C.R., McIntyre, L., Earl, E.A., Klein, R.L., Shimony, J.S., Petersen, S.E., Schlaggar, B.L., Fair, D.A., Dosenbach, N.U.F., 2018. Behavioral interventions for reducing head motion during MRI scans in children. Neuroimage. https://doi.org/10.1016/j.neuroimage.2018.01.023

Greve, D.N., Fischl, B., 2009. Accurate and robust brain image alignment using boundary-based registration. Neuroimage 48, 63–72. https://doi.org/10.1016/j.neuroimage.2009.06.060

Griffanti, L., Salimi-Khorshidi, G., Beckmann, C.F., Auerbach, E.J., Douaud, G., Sexton, C.E., Zsoldos, E., Ebmeier, K.P., Filippini, N., Mackay, C.E., Moeller, S., Xu, J., Yacoub, E., Baselli, G., Ugurbil, K., Miller, K.L., Smith, S.M., 2014. ICA-based artefact removal and accelerated fMRI acquisition for improved resting state network imaging. Neuroimage 95, 232–247. https://doi.org/10.1016/j.neuroimage.2014.03.034

Hallquist, M.N., Hwang, K., Luna, B., 2013. The nuisance of nuisance regression: Spectral misspecification in a common approach to resting-state fMRI preprocessing reintroduces noise and obscures functional connectivity. Neuroimage 82C, 208–225. https://doi.org/10.1016/j.neuroimage.2013.05.116

Jenkinson, M., Beckmann, C.F., Behrens, T.E.J., Woolrich, M.W., Smith, S.M., 2012. FSL. Neuroimage 62, 782–790. https://doi.org/10.1016/j.neuroimage.2011.09.015

Jo, H.J., Gotts, S.J., Reynolds, R.C., Bandettini, P.A., Martin, A., Cox, R.W., Saad, Z.S., 2013. Effective preprocessing procedures virtually eliminate distance-dependent motion artifacts in resting state FMRI. J. Appl. Math. 2013. https://doi.org/10.1155/2013/935154

Karalunas, S.L., Fair, D., Musser, E.D., Aykes, K., Iyer, S.P., Nigg, J.T., 2014. Subtyping attention-deficit/hyperactivity disorder using temperament dimensions: toward biologically based nosologic criteria. JAMA psychiatry 71, 1015–24. https://doi.org/10.1001/jamapsychiatry.2014.763

Karalunas, S.L., Geurts, H.M., Konrad, K., Bender, S., Nigg, J.T., 2014. Annual research review: Reaction time variability in ADHD and autism spectrum disorders: Measurement and mechanisms of a proposed trans-diagnostic phenotype. J. Child Psychol. Psychiatry Allied Discip. https://doi.org/10.1111/jcpp.12217

Kundu, P., Brenowitz, N.D., Voon, V., Worbe, Y., Vertes, P.E., Inati, S.J., Saad, Z.S., Bandettini, P.A., Bullmore, E.T., 2013. Integrated strategy for improving functional connectivity mapping using multiecho fMRI. Proc. Natl. Acad. Sci. 110, 16187–16192. https://doi.org/10.1073/pnas.1301725110

Laumann, T.O., Snyder, A.Z., Mitra, A., Gordon, E.M., Gratton, C., Adeyemo, B., Gilmore, A.W., Nelson, S.M., Berg, J.J., Greene, D.J., McCarthy, J.E., Tagliazucchi, E., Laufs, H., Schlaggar, B.L., Dosenbach, N.U.F., Petersen, S.E., 2016. On the Stability of BOLD fMRI Correlations. Cereb. Cortex 1–14. https://doi.org/10.1093/cercor/bhw265

Lisdahl, K.M., Sher, K.J., Conway, K.P., Gonzalez, R., Feldstein Ewing, S.W., Nixon, S.J., Tapert, S., Bartsch, H., Goldstein, R.Z., Heitzeg, M., 2018. Adolescent brain cognitive development (ABCD) study: Overview of substance use assessment methods. Dev. Cogn. Neurosci. https://doi.org/10.1016/j.dcn.2018.02.007

Mills, B.D., Miranda-Dominguez, O., Mills, K., Earl, E., Cordova, M., Painter, J., Karalunas, S.L., Nigg, J.T., Fair, D.A., 2017. ADHD and Attentional Control: Impaired Segregation of Task Positive and Task Negative Brain Networks. Netw. Neurosci. 1–36. https://doi.org/10.1162/NETN_a_00034

Mills, K.L., Bathula, D., Dias, T.G.C., Iyer, S.P., Fenesy, M.C., Musser, E.D., Stevens, C.A., Thurlow, B.L., Carpenter, S.D., Nagel, B.J., Nigg, J.T., Fair, D.A., 2012. Altered cortico-striatal-thalamic connectivity in relation to spatial working memory capacity in children with ADHD. Front Psychiatry 3, 2. https://doi.org/10.3389/fpsyt.2012.00002

Mills, K.L., Bathula, D., Dias, T.G.C., Iyer, S.P., Fenesy, M.C., Musser, E.D., Stevens, C.A., Thurlow, B.L., Carpenter, S.D., Nagel, B.J., Nigg, J.T., Fair, D.A., 2012. Altered Cortico-Striatal-Thalamic Connectivity in Relation to Spatial Working Memory Capacity in Children with ADHD. Front. Psychiatry 3. https://doi.org/10.3389/fpsyt.2012.00002

Miranda-Dominguez, O., Feczko, E., Grayson, D.S.D.S., Wallum, H., Nigg, J.T.J.T., Fair, D.A.D.A., 2017. Heritability of the human connectome: a connectotyping study. Netw. Neurosci. 1–48. https://doi.org/10.1162/NETN_a_00029

Moeller, S., Yacoub, E., Olman, C.A., Auerbach, E., Strupp, J., Harel, N., Uğurbil, K., 2010. Multiband multislice GE-EPI at 7 tesla, with 16-fold acceleration using partial parallel imaging with application to high spatial and temporal whole-brain FMRI. Magn. Reson. Med. 63, 1144–1153. https://doi.org/10.1002/mrm.22361

Muschelli, J., Nebel, M.B., Caffo, B.S., Barber, A.D., Pekar, J.J., Mostofsky, S.H., 2014. Reduction of motion-related artifacts in resting state fMRI using aCompCor. Neuroimage 96, 22–35. https://doi.org/10.1016/j.neuroimage.2014.03.028

Nigg, J.T., Gustafsson, H.C., Karalunas, S.L., Ryabinin, P., McWeeney, S., Faraone, S. V., Mooney, M., Fair, D.A., Wilmot, B., 2018. Working Memory and Vigilance as Multivariate Endophenotypes Related to Common Genetic Risk for Attention-Deficit/Hyperactivity Disorder. J. Am. Acad. Child Adolesc. Psychiatry 57, 175–182. https://doi.org/10.1016/j.jaac.2017.12.013

Patel, A.X., Kundu, P., Rubinov, M., Jones, P.S., Vértes, P.E., Ersche, K.D., Suckling, J., Bullmore, E.T., 2014. A wavelet method for modeling and despiking motion artifacts from resting-state fMRI time series. Neuroimage 95, 287–304. https://doi.org/10.1016/j.neuroimage.2014.03.012

Power, J., Barnes, K., Snyder, A., Schlaggar, B., Petersen, S., 2013. Steps toward optimizing motion artifact removal in functional connectivity MRI; a reply to Carp. Neuroimage 76, 439–441. https://doi.org/10.1016/j.neuroimage.2012.03.017

Power, J., Barnes, K., Snyder, A., Schlaggar, B., Petersen, S., 2012. Spurious but systematic correlations in functional connectivity MRI networks arise from subject motion. Neuroimage 59, 2142–2154. https://doi.org/10.1016/j.neuroimage.2011.10.018

Power, J., Mitra, A., Laumann, T., Snyder, A., Schlaggar, B., Petersen, S., 2014. Methods to detect, characterize, and remove motion artifact in resting state fMRI. Neuroimage 84, 320–341. https://doi.org/10.1016/j.neuroimage.2013.08.048

Power, J.D., 2016. A simple but useful way to assess fMRI scan qualities. Neuroimage. https://doi.org/10.1016/j.neuroimage.2016.08.009

Power, J.D., Plitt, M., Laumann, T.O., Martin, A., 2017. Sources and implications of whole-brain fMRI signals in humans. Neuroimage 146, 609–625. https://doi.org/10.1016/j.neuroimage.2016.09.038

Pruim, R.H.R., Mennes, M., van Rooij, D., Llera, A., Buitelaar, J.K., Beckmann, C.F., 2015. ICA-AROMA: A robust ICA-based strategy for removing motion artifacts from fMRI data. Neuroimage 112, 267–277. https://doi.org/10.1016/j.neuroimage.2015.02.064

Ray, S., Miller, M., Karalunas, S., Robertson, C., Grayson, D.S., Cary, R.P., Hawkey, E., Painter, J.G., Kriz, D., Fombonne, E., Nigg, J.T., Fair, D.A., 2014. Structural and functional connectivity of the human brain in autism spectrum disorders and attention-deficit/hyperactivity disorder: A rich club-organization study. Hum. Brain Mapp. https://doi.org/10.1002/hbm.22603

Salimi-Khorshidi, G., Douaud, G., Beckmann, C.F., Glasser, M.F., Griffanti, L., Smith, S.M., 2014. Automatic denoising of functional MRI data: Combining independent component analysis and hierarchical fusion of classifiers. Neuroimage 90, 449–468. https://doi.org/10.1016/j.neuroimage.2013.11.046

Satterthwaite, T., Wolf, D., Loughead, J., Ruparel, K., Elliott, M., Hakonarson, H., Gur, R., Gur, R., 2012. Impact of in-scanner head motion on multiple measures of functional connectivity: relevance for studies of neurodevelopment in youth. Neuroimage 60, 623–632. https://doi.org/10.1016/j.neuroimage.2011.12.063

Satterthwaite, T.D., Elliott, M.A., Gerraty, R.T., Ruparel, K., Loughead, J., Calkins, M.E., Eickhoff, S.B., Hakonarson, H., Gur, R.C., Gur, R.E., Wolf, D.H., 2013. An improved framework for confound regression and filtering for control of motion artifact in the preprocessing of resting-state functional connectivity data. Neuroimage 64, 240–256. https://doi.org/10.1016/j.neuroimage.2012.08.052

Siegel, J.S., Mitra, A., Laumann, T.O., Seitzman, B.A., Raichle, M., Corbetta, M., Snyder, A.Z., 2016. Data Quality Influences Observed Links Between Functional Connectivity and Behavior. Cereb. Cortex 1–11. https://doi.org/10.1093/cercor/bhw253

Siegel, J.S., Mitra, A., Laumann, T.O., Seitzman, B.A., Raichle, M., Corbetta, M., Snyder, A.Z., 2016. Data Quality Influences Observed Links Between Functional Connectivity and Behavior. Cereb. Cortex. https://doi.org/10.1093/cercor/bhw253

Smith, S.M., Jenkinson, M., Woolrich, M.W., Beckmann, C.F., Behrens, T.E.J., Johansen-Berg, H., Bannister, P.R., De Luca, M., Drobnjak, I., Flitney, D.E., Niazy, R.K., Saunders, J., Vickers, J., Zhang, Y., De Stefano, N., Brady, J.M., Matthews, P.M., 2004. Advances in functional and structural MR image analysis and implementation as FSL. Neuroimage 23 Suppl 1, S208–19. https://doi.org/10.1016/j.neuroimage.2004.07.051

Smith, S.M., Jenkinson, M., Woolrich, M.W., Beckmann, C.F., Behrens, T.E.J., Johansen-Berg, H., Bannister, P.R., De Luca, M., Drobnjak, I., Flitney, D.E., Niazy, R.K., Saunders, J., Vickers, J., Zhang, Y., De Stefano, N., Brady, J.M., Matthews, P.M., 2004. Advances in functional and structural MR image analysis and implementation as FSL. Neuroimage 23, S208–S219. https://doi.org/10.1016/j.neuroimage.2004.07.051

Smyser, C.D., Inder, T.E., Shimony, J.S., Hill, J.E., Degnan, A.J., Snyder, A.Z., Neil, J.J., 2010. Longitudinal analysis of neural network development in preterm infants. Cereb. Cortex 20, 2852–62. https://doi.org/10.1093/cercor/bhq035

Tisdall, M.D., Reuter, M., Qureshi, A., Buckner, R.L., Fischl, B., van der Kouwe, A.J.W., 2016. Prospective motion correction with volumetric navigators (vNavs) reduces the bias and variance in brain morphometry induced by subject motion. Neuroimage 127, 11–22. https://doi.org/10.1016/j.neuroimage.2015.11.054

Todd, N., Moeller, S., Auerbach, E.J., Yacoub, E., Flandin, G., Weiskopf, N., 2016. Evaluation of 2D multiband EPI imaging for high-resolution, whole-brain, task-based fMRI studies at 3T: Sensitivity and slice leakage artifacts. Neuroimage 124, 32–42. https://doi.org/10.1016/j.neuroimage.2015.08.056

Uğurbil, K., Xu, J., Auerbach, E.J., Moeller, S., Vu, A.T., Duarte-Carvajalino, J.M., Lenglet, C., Wu, X., Schmitter, S., Van de Moortele, P.F., Strupp, J., Sapiro, G., De Martino, F., Wang, D., Harel, N., Garwood, M., Chen, L., Feinberg, D. a., Smith, S.M., Miller, K.L., Sotiropoulos, S.N., Jbabdi, S., Andersson, J.L.R., Behrens, T.E.J., Glasser, M.F., Van Essen, D.C., Yacoub, E., WU-Minn HCP Consortium, 2013. Pushing spatial and temporal resolution for functional and diffusion MRI in the Human Connectome Project. Neuroimage 80, 80–104. https://doi.org/10.1016/j.neuroimage.2013.05.012

Van de Moortele, P., Pfeuffer, J., Glover, G.H., Ugurbil, K., Hu, X., 2002. Respiration-induced B0 fluctuations and their spatial distribution in the human brain at 7 Tesla. Magn Reson Med 47, 888–895. https://doi.org/10.1002/mrm.10145

Van Dijk, K.R. a, Sabuncu, M.R., Buckner, R.L., 2012. The influence of head motion on intrinsic functional connectivity MRI. Neuroimage 59, 431–438. https://doi.org/10.1016/j.neuroimage.2011.07.044

Van Dijk, K.R., Sabuncu, M.R., Buckner, R.L., 2012. The influence of head motion on intrinsic functional connectivity MRI. Neuroimage 59, 431–438. https://doi.org/10.1016/j.neuroimage.2011.07.044

Volkow, N.D., Koob, G.F., Croyle, R.T., Bianchi, D.W., Gordon, J.A., Koroshetz, W.J., Pérez-Stable, E.J., Riley, W.T., Bloch, M.H., Conway, K., Deesds, B.G., Dowling, G.J., Grant, S., Howlett, K.D., Matochik, J.A., Morgan, G.D., Murray, M.M., Noronha, A., Spong, C.Y., Wargo, E.M., Warren, K.R., Weiss, S.R.B., 2017. The conception of the ABCD study: From substance use to a broad NIH collaboration. Dev. Cogn. Neurosci. 1–4. https://doi.org/10.1016/j.dcn.2017.10.002

Wallis, L.A., Healy, M., Undy, M.B., Maconochie, I., 2005. Age related reference ranges for respiration rate and heart rate from 4 to 16 years. Arch. Dis. Child. 90, 1117–1121. https://doi.org/10.1136/adc.2004.068718

Woolrich, M.W., Jbabdi, S., Patenaude, B., Chappell, M., Makni, S., Behrens, T., Beckmann, C., Jenkinson, M., Smith, S.M., 2009. Bayesian analysis of neuroimaging data in FSL. Neuroimage 45, S173–86. https://doi.org/10.1016/j.neuroimage.2008.10.055

Xu, J., Moeller, S., Auerbach, E.J., Strupp, J., Smith, S.M., Feinberg, D.A., Yacoub, E., Uğurbil, K., 2013. Evaluation of slice accelerations using multiband echo planar imaging at 3T. Neuroimage 83, 991–1001. https://doi.org/10.1016/j.neuroimage.2013.07.055

Yan, C.G., Cheung, B., Kelly, C., Colcombe, S., Craddock, R.C., Di Martino, A., Li, Q., Zuo, X.N., Castellanos, F.X., Milham, M.P., 2013. A comprehensive assessment of regional variation in the impact of head micromovements on functional connectomics. Neuroimage 76, 183–201. https://doi.org/10.1016/j.neuroimage.2013.03.004

Zaitsev, M., Akin, B., LeVan, P., Knowles, B.R., 2017. Prospective motion correction in functional MRI. Neuroimage 154, 33–42. https://doi.org/10.1016/j.neuroimage.2016.11.014

